# *Bacteroides uniformis* enhances endurance exercise performance through gluconeogenesis

**DOI:** 10.1101/2020.03.04.975730

**Authors:** Hiroto Morita, Chie Kano, Chiharu Ishii, Noriko Kagata, Takamasa Ishikawa, Yoshihide Uchiyama, Susumu Hara, Teppei Nakamura, Shinji Fukuda

**Author notes:** H.M. and C.K. contributed equally to this work. Corresponding author: Shinji Fukuda, Institute for Advanced Biosciences, Keio University, 246-2 Mizukami, Kakuganji, Tsuruoka, Yamagata 997-0052, Japan.

## Abstract

Athletes require high levels of energy to exercise under extreme conditions. Gut microbiota supplies energy to the host; however, the mechanism how gut microbiota contribute in the athlete is unclear. In this study, we determined that gut microbiota of Japanese long-distance runners differed from that of non-athletes, and the *Bacteroides uniformis* cell number in the feces correlated with 3,000-m race time. Mice administrated with *B. unformis* extended the swimming time to exhaustion. Furthermore, acetate and propionate concentrations in the cecum increased in *B. uniformis*-administered mice subjected to weekly exercise. Expression levels of carnitine palmitoyl transferase 1a and phosphoenolpyruvate carboxykinase genes were elevated in the liver, suggesting that acetate and propionate produced by *B. uniformis* improve endurance exercise performance, at least in part, through enhancing gluconeogenesis. In addition, α-cyclodextrin administration increased *B. uniformis* and improved the performance in humans and mice, thus it is a candidate substance enhancing exercise performance through modification of gut microbiota.

## INTRODUCTION

Human exercise performance comprises a combination of physical strength (e.g., muscle and explosive strength, endurance, and flexibility) and the actions performed in a given exercise (e.g., throwing, kicking, and hitting). Exercise performance can be improved through training; however, other factors, such as genetic background, may affect it. Recently, several reports have demonstrated that the gut microbiota contributes to health, energy homeostasis, and glucose and lipid metabolisms ^1–4^. Hsu *et al*. reported that the swimming endurance time of germ-free mice was shorter than that of specific-pathogen-free mice and *Bacteroides fragilis*-monoassociated mice, thus identifying gut microbiota as one of the factors associated with exercise performance ^5^. Conversely, other studies have shown that exercise changes the gut microbiome profile in mice ^6,7^. In humans, Clarke *et al*. reported that the gut microbiota of rugby players had high α diversity and relative abundance of the genera *Prevotella, Succinivibrio, Bacteroides*, and *Lactobacillus*, as a difference to those of non-rugby players ^8^. Another study on the gut microbiota of cyclists showed that a high abundance of *Prevotella* was correlated with weekly exercise time ^9^. However, none of these studies demonstrated a causal relationship between exercise performance and characteristic genera in the gut microbiota of athletes. Recently, *Veillonella atypica*, isolated from the gut microbiota of marathon runners, was shown to produce propionate from exercise-induced lactate in the gut, improving endurance exercise performance in mice ^10^ Only that report has demonstrated a causal relationship between exercise performances and the gut microbiota in mice, but detailed relationship remains unclear. In the present study, we investigated the role of *Bacteroides uniformis*,identified in the microbiome analysis of Japanese long-distance runners, in the endurance exercise. We observed that administration of *B. uniformis* in mice improved their endurance exercise performance. We also evidenced elevated transcriptional levels of *Cpt1a* (encoding the carnitine palmitoyl transferase 1a that regulates βoxidation of fatty acids) and *Pck1* (encoding the phosphoenolpyruvate carboxykinase that regulates gluconeogenesis) in the liver of mice administered with *B. uniformis* and subjected to exercise. The concentration of short-chain fatty acids (SCFAs), such as acetate and propionate, in the cecum of the mice was higher than that of control mice, suggesting transcriptional changes of the genes were due to SCFAs produced by *B. uniformis*. In addition, α-cyclodextrin, which increased the cell number of *B. uniformis* in human and murine intestine, improved endurance exercise performance, and reduced post-exercise fatigue in a human study. Our findings may contribute to a better understanding on how the gut microbiota relates to an increased endurance exercise performance and how the energy required for exercise is obtained.

## RESULTS

### Gut microbiome profile differed between Japanese long-distance runners and non-athletes

We collected faecal samples from 48 male students (age 20.3 ± 1.0 years [mean ± s.d]; height 169.9 ± 4.2 cm; body weight 54.8 ± 3.5 kg; and body mass index (BMI) 19.0 ± 0.9), who were members of the Aoyama Gakuin University Track and Field Club Ekiden team in July 2015 (athlete group). Faecal samples were also taken from ten other men in their twenties (age 22.6 ± 0.7 years; height 169.7 ± 7.6 cm; body weight 59.5 ± 8.0 kg; BMI 20.6 ± 2.2) as controls (non-athlete group). DNA was extracted from the feces for sequencing of the V4 region of the bacterial 16S rRNA gene with the MiSeq system to obtain information on gut microbiome profile. Five subjects in the athlete group and two subjects in the non-athlete group were excluded from analysis because the polymerase chain reaction (PCR) failed to amplify the sequence when MiSeq libraries were constructed. The data revealed that α diversity was significantly greater in the athlete group than in the non-athlete group according to Chao1, the Shannon index, and phylogenetic distance (PD) whole tree (Fig. 1a). Analysis of β diversity based on weighted and unweighted UniFrac distances also revealed a significantly different microbiome profile between the two groups (Fig. 1b). The LEfSe analysis indicated differences in microbiome profile, specifically in the following 9 genera: *Bacteroides, Prevotella, Lachnospira, Sutterella*, an unknown genus of the family Gemellaceae, *Granulicatella*, *Escherichia*, *Citrullus*, and an unknown genus of the order *Streptophyta* (Fig. 1c). Comparison of the relative abundance of these 9 genera between groups revealed significant differences in six of them: *Bacteroides, Lachnospira, Sutterella, Escherichia*, Gemellaceae, and *Streptophyta* (Supplementary Table 1).

**Fig. 1:**
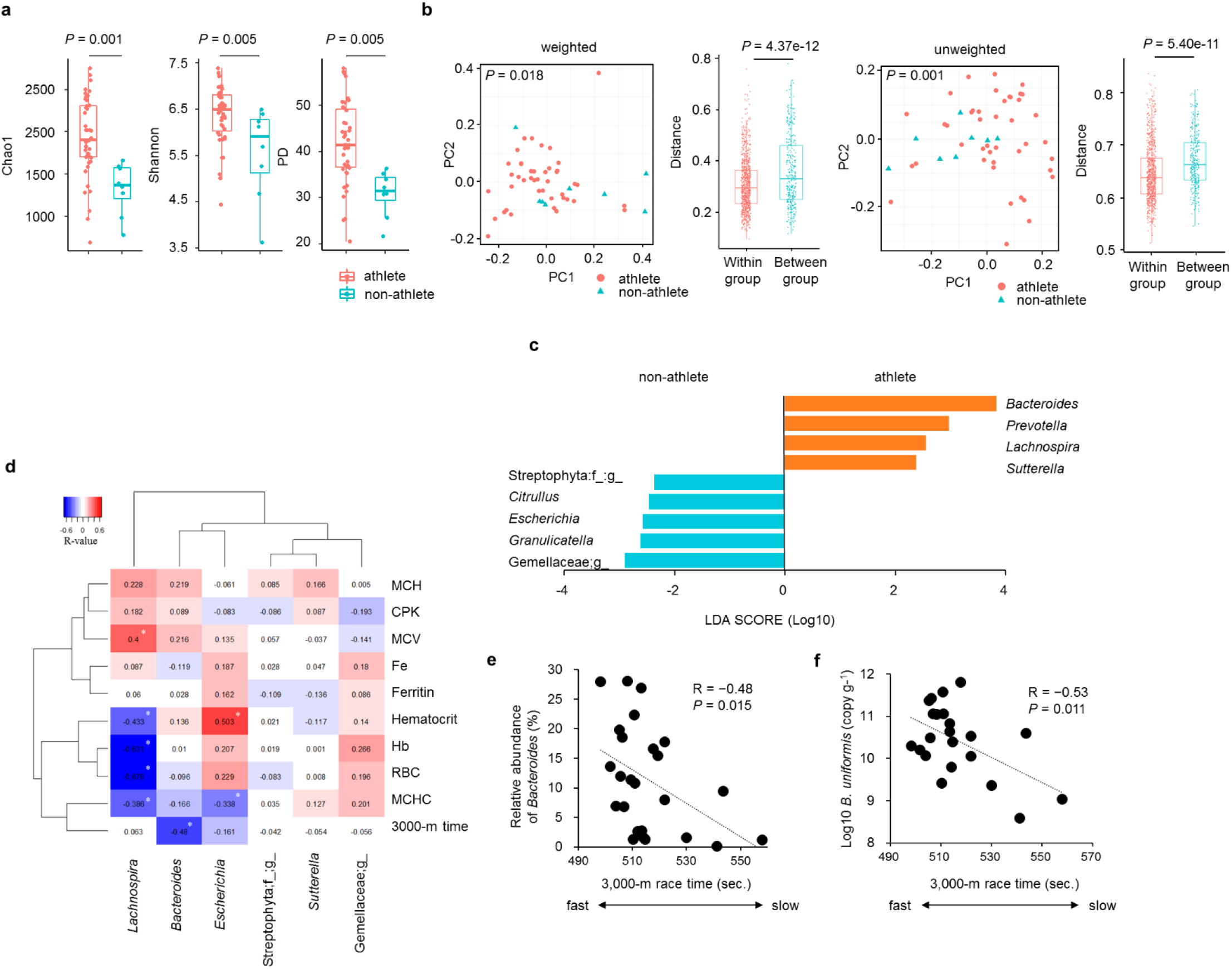
Abundance of *Bacteroides* is higher in athletes and it correlates with race time. a, Comparison of microbiota α diversity between athletes and non-athletes. Data are presented as median with first and third quartiles represented by the box edges. The whiskers correspond to the minimum and maximum values. Statistical significance of differences between groups was analysed by a two-sample *t*-test using Monte Carlo permutations. b, Comparison of microbiota β diversity between athletes and non-athletes based on weighted and unweighted UniFrac distance. Statistical significance of differences between groups was analysed by the PREMANOVA. c, The 9 genera of bacteria most likely explained the differences between the microbiota of athletes and non-athletes. Analysis was conducted with LEfSe to abstract genera with an | LDA score | > 2. The bars represent the LDA score. d, Analysis of correlations of3,000-m race time and blood parameters with the relative abundance of the 6 genera in the athlete group. The genera were selected based on | LDA score | > 2 in LEfSe and significant difference in relative abundance. Values in the cells represent Spearman’s rank correlation coefficients. e, Scatterplot of the relative abundance of *Bacteroides* in athlete group and 3,000-m race time. Correlation was determined using the Spearman rank correlation coefficient. Dotted line shows the approximated curve. f, Scatterplot of log10 cell number of *B. uniformis* in athletes and 3,000-m race time. Bacterial count was based on 16S rRNA gene copy number that was calculated by qPCR. Correlation was determined using the Pearson correlation coefficient. Dotted line shows the approximated curve. Athlete group, *n* = 43; non-athlete group *n* = 8 for all parameters, except for 3,000-m race time: athlete group, *n* = 25. Eighteen of 43 athletes refused to measure the time. All tests were two-tailed. * *P* < 0.05. PREMANOVA: permutational multivariate analysis of variance; LDA: linear discriminant analysis; MCH: mean corpuscular hemoglobin; CPK: creatine phosphokinase; MCV: mean corpuscular volume; Hb: hemoglobin; RBC: red blood cells; MCHC: mean corpuscular hemoglobin concentration.

To search the association between gut microbiota and endurance exercise performance, we analysed the correlations of the relative abundance of each of the above six genera with blood parameters and 3000-m race time. The blood parameters were red blood cell count (RBC), mean corpuscular volume (MCV), mean corpuscular hemoglobin (MCH), mean corpuscular hemoglobin concentration (MCHC), hemoglobin concentration (Hb), hematocrit, serum iron (Fe), and ferritin levels, all of which are associated with anemia and may affect long-distance running capacity; and creatin phosphokinase (CPK) levels, which reflect the degree of muscle injury. This analysis showed that only the abundance of *Bacteroides* was significantly and negatively correlated with 3000-m race time (Fig. 1d and e).

Next, to examine which species of *Bacteroides* were associated with endurance exercise performance, we calculated the cell number of thirteen species commonly detected in human feces by using species-specific quantitative PCR. As shown in Supplementary Table 2, *Bacteroides caccae, Bacteroides eggerthii*, *B. uniformis*, *Bacteroides thetaiotaomicron*, *Bacteroides vulgatus*, and *Bacteroides dorei* were more significantly abundant in the athlete group than in the non-athlete group. Analyses of the cell numbers of these 6 species with 3000-m race time revealed a significant negative correlation between 3000-m race time and *B. uniformis* cell number (Fig. 1f).

### Administration of *B. uniformis* prolonged endurance exercise and altered intestinal SCFAs and bile acid composition

To determine the relationship between *B. uniformis* and endurance exercise performance, we administered mice with *B. uniformis* orally and conducted a weekly swimming endurance test (Fig. 2a). Amazingly, at 4 weeks, mice treated with *B. uniformis* could swim for significantly longer time than a control group administered with phosphate-buffered saline (PBS) (Fig. 2b). This result indicated that the administration of *B. uniformis* improved endurance performance in mice.

**Fig. 2:**
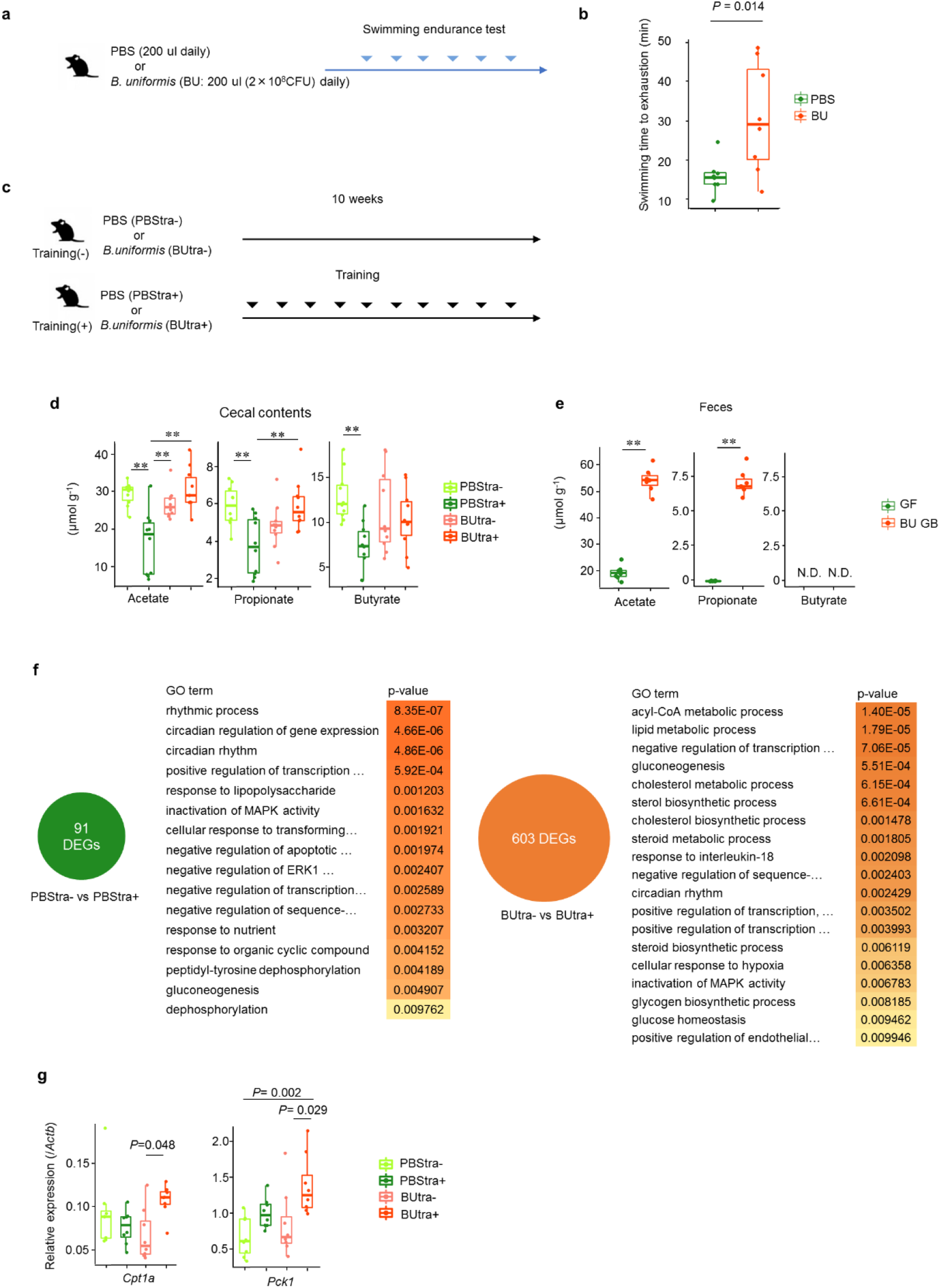
*B. uniformis* administration improved endurance exercise performance and altered the intestinal SCFAs. a, Endurance test schedule. Mice were administered phosphate-bufered saline (PBS) or *B. uniformis* daily and subjected to swimming endurance test weekly b, Swimming time to exhaustion of PBS administrated mice (*n* = 8) and *B. uniformis* administrated mice (*n* = 8) on week 4. c, Experiment schedule. Mice were administered PBS or *B. uniformis* daily. The training (+) groups were subjected to weekly swimming exercise. d, Concentrations of SCFAs in cecal content ofPBStra− (*n* = 10), PBStra+ (*n* = 10), BUtra− (*n* = 10), and BUtra+ (*n* = 10). e, Concentrations of SCFAs in feces of germ-free mice (*n* = 8) and *B. uniformis-monoassociated* mice (*n* = 7). f, Gene Ontology (GO) analysis conducted for genes that demonstrated different expression levels in the liver. Differences in expression were observed for 91 genes in PBStra− vs PBStra+ and in 603 genes in BUtra− vs Butra+. These genes are related to the GO terms shown on the right side of the panel. GO analysis were carried out using DAVID online tool. Statistical significance were analysed by Fisher’s exact test. *P*<0.01 was considered significant. g, Comparisons of hepatic expression of *Cpt1a* and *Pck1* in PBStra− (*n* = 8), PBStra+ (*n* = 8), BUtra− (*n* = 8), and BUtra+ (*n* = 8) measured by RT-qPCR. Data are presented as median with first and third quartiles as the box edges. The whiskers correspond to the minimum and maximum. Statistical significance of difference between groups were analysed by Mann–Whitney *t*-test (b), Tukey-Kramer test (d,g), unpaired *t*-test (e). All tests were two-tailed. ** *P* < 0.01 BU: mice administered *B. uniformis*. PBStra+: mice administered PBS and subjected to exercise; PBStra−: mice administered PBS and not subjected to exercise; BUtra+: mice administered *B. uniformis* and subjected to exercise; BUtra−: mice administered *B. uniformis* and not subjected to exercise; GF: germ-free mice; GB: gnotobiotic mice monoassociated with *B. uniformis* type strain; N.D.: not detectable; DEG: differentially expressed gene.

Next, mice were forced to swim for 15 min once a week (Fig. 2c) but in milder conditions than those used in the swimming endurance test to induce uniform response of the mice to exercise. Immediately after the swimming exercise on day 10, mice were sacrificed, the cecal contents were collected, and SCFAs concentrations were measured. The concentrations of acetate, propionate, and butyrate were all significantly lower in mice administered PBS with exercise (PBStra+), compared to mice administered PBS without exercise (PBStra−) (Fig. 2d). However, mice administered *B. uniformis* with exercise (BUtra+) exhibited similar concentrations of the above three SCFAs than those of mice administered *B. uniformis* without exercise (BUtra−), and the concentrations of acetate and propionate were significantly higher compared with those in PBStra+ (Fig. 2d). These results indicated that while exercise reduced concentrations of SCFAs in cecum, the concentrations were not decreased when *B. uniformis* was administered. These facts implied that the presence of *B. uniformis* in the gut inhibited absorption of the SCFAs from cecum and/or produced SCFAs at a higher rate than their absorption rate. Therefore, we next used *B. uniformis-monoassociated* mice to examine whether *B. uniformis* produced these SCFAs in the murine intestine. We evidenced that *B. uniformis* produced acetate and propionate in the murine intestine. (Fig. 2e).

Since SCFAs are known as energy substrates for the colonic epithelium and peripheral tissues ^11^, we performed RNA-seq analysis of mRNAs extracted from liver, and conducted gene ontology (GO) analysis of differentially expressed genes with a *q*-value < 0.05. GO analysis when comparing PBStra− and PBStra+ groups revealed enrichment in rhythmic process and circadian rhythm (Fig. 2f). It was previously reported that the expression of transcriptional repressor *Per2*, a component of the circadian clock, was modified by exercise during light phase ^12,13^. Similarly, *Per2* expression increased in the PBStra+ group in our experiment. On the other hand, comparison between BUtra− and BUtra+ groups indicated enrichment of lipid metabolism and gluconeogenesis in addition to GO terms related to circadian rhythm (Fig. 2f). Specifically, compared to BUtra− mice, BUtra+ mice liver showed elevated expression of *Cpt1a* (GO term: lipid metabolism) and *Pck1* (GO term: gluconeogenesis) (Fig. 2g). *Cpt1a* codes for carnitine palmitoyl transferase 1a that transfers long chain fatty acids into mitochondria for β oxidation ^14^ Meanwhile, *Pck1* codes for phosphoenolpyruvate carboxykinase that catalyses the reversible conversion of oxaloacetate to phosphoenolpyruvate for gluconeogenesis ^15^.

Bile acids not only aid absorption of dietary lipids but also function as signalling factor controlling glucose, lipid, and energy metabolism in multiple organs, such as gut, brown adipose tissue, and skeletal muscle ^16^ As shown in Supplementary Fig. 1a, while no significant differences in total bile acid concentration in the liver were noted, levels of glycocholic acid (GCA), were significantly lower in PBStra+ than in PBStra− group; however, no significant reduction was observed in comparison to levels of BUtra+ and BUtra− groups. While levels of β-muricholic acid (β-MCA), the main primary bile acid in mice, did not significantly differ between PBStra+ and PBStra− groups, levels were significantly higher in BUtra+ than in BUtra−. Similarly, levels of deoxycholic acid (DCA), the secondary bile acids, did not significantly differ between PBStra+ and PBStra− groups but were significantly higher in BUtra+ than in BUtra−. Concentrations of hyodeoxycholic acid (HDCA), α-muricholic acid (α-MCA) and ω-muricholic acid (ω-MCA) were significantly higher in BUtra+ group, compared to those present in the other three groups. These results suggested that the bile acid composition in the liver differed between the control and *B. uniformis-treated* group even under the same exercise condition. Especially, the bile acid composition of BUtra+ group had elevated levels of MCA and its secondary bile acids. However, *B. uniformis* inoculation did not affect the expression of the gene encoding Cyp2c70, which converts chenodeoxycholic acid (CDCA) into α-MCA and β-MCA, or any other genes associated with bile acid production (Supplementary Fig. 1b).

### Administration of α-cyclodextrin increased *B. uniformis* cell number in the murine intestine and improved endurance exercise performance

*B. uniformis* is one of the dominant bacteria in the human gut microbiota ^17^, but its ingestion has not been studied, specifically in terms of safety in humans. Therefore, to explore the application of the above findings, we searched for food that specifically favoured proliferation of *B. uniformis*, such as secoisolariciresinol diglucoside (SDG), an antioxidant phytoestrogen found in flaxseed and other seeds. Interestingly, *B. uniformis* expresses a gene that encodes β-glucosidase that releases glucose from SDG via hydrolysis ^18,19^. We therefore feed mice with flaxseed lignans, which are high in SDG, and examined the proliferation of *B. uniformis* in the gut. The food-grade flaxseed lignans used in this experiment contained approximately 40% (w w^-1^) α-cyclodextrin as an inclusion compound. Alpha-cyclodextrin, a member of the cyclic oligosaccharide family composed of six glucose subunits, was reported to change the gut microbiota and reduce weight gain of epididymal adipose tissue in mice fed with a high-fat diet ^20^ We therefore similarly examined the activity of α-cyclodextrin. Mice were provided with standard CE-2 diet containing flaxseed lignans, *ad libitum* (FL group) (Fig. 3a). FL group evidenced a significantly higher relative abundance of genus *Bacteroides* and cell number of *B. uniformis* in feces than a control group at 7 weeks (Fig. 3b, c). Furthermore, the FL group exhibited a significantly longer swimming time to exhaustion in the swimming endurance test (Fig. 3d). Importantly, a similar and significant increase in *B. uniformis* cell number in feces and an extension in swimming time to exhaustion was noted in mice treated with α-cyclodextrin diet (αCD group) (Fig. 3e-h).

**Fig. 3:**
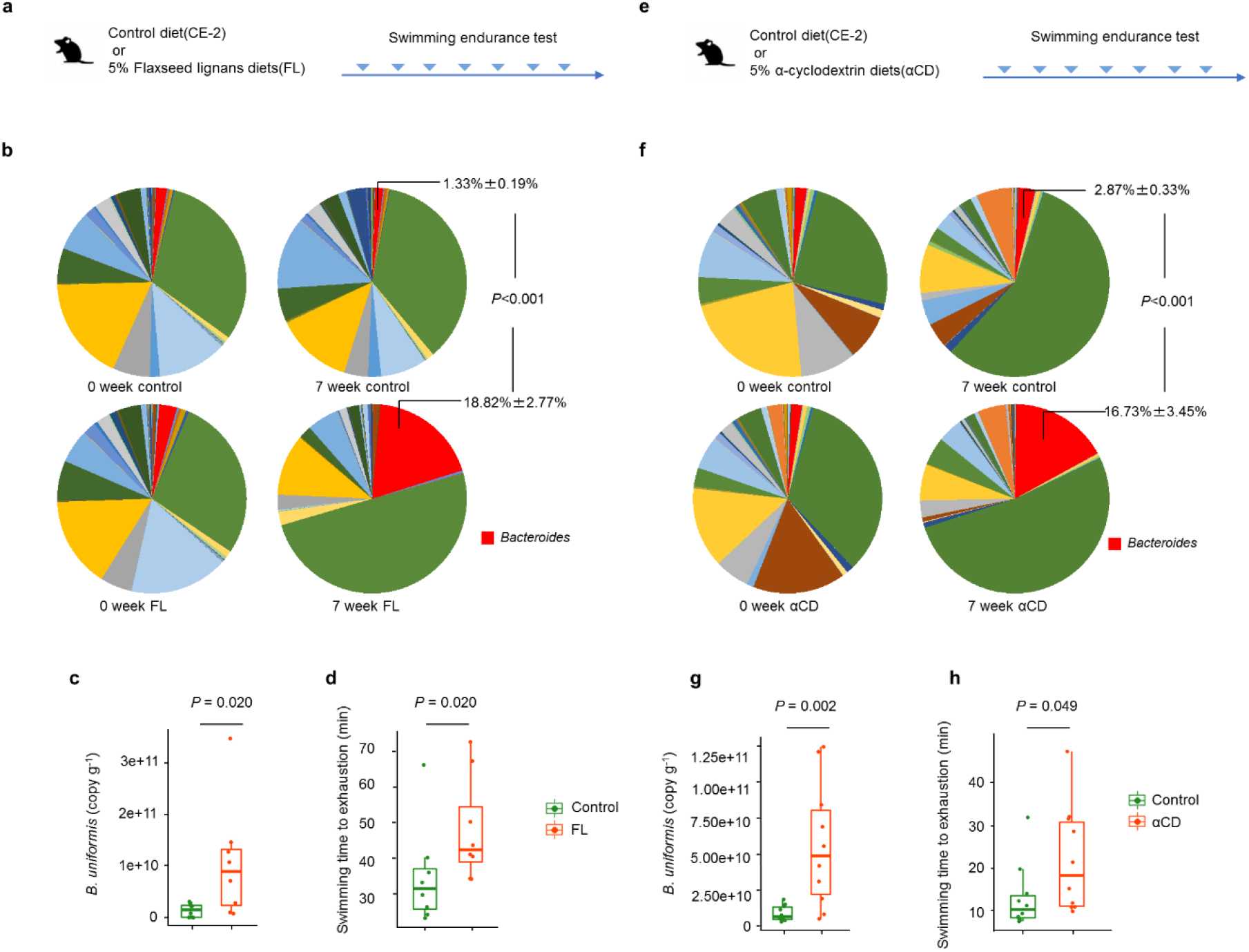
Flaxseed lignans (FL) and α-cyclodextrin (αCD) increased the cell number of *B. uniformis* in the murine intestine and prolonged their swimming time to exhaustion. a,e, Endurance test schedule. Mice were fed standard diet containing FL or αCD and subjected to weekly swimming endurance test (*n* = 8 for FL group and control for FL group, *n* = 10 for αCD group and control for αCD group). b,f, Relative abundance of *Bacteroids* (mean ±s.e.m.) in FL group (b) and αCD group (f). Each area in pie charts shows the mean relative abundance of each bacterial genera. c,g, Cell number of *B. uniformis* in feces from each group at 7 weeks. Bacterial count was based on 16S rRNA gene copy number that was calculated by species-specific qPCR. d,h, Swimming time to exhaustion of each group at 7 weeks. Data are presented as median with first and third quartiles as the box edges, and the whiskers correspond to the minimum and maximum (c,d,g,h). Statistical significance of difference between groups were analysed by two-tailed Mann-Whitney *U*-tests.

### Alpha-cyclodextrin increased the *B. uniformis* cell number and exercise performance in humans

To confirm the effects of flaxseed lignans and α-cyclodextrin on human exercise, we conducted a randomised, double-blind, placebo-controlled, parallel-group study in humans. Thirty-six Japanese men who had a habit to exercise (once or twice exercise per week) were randomly assigned to three different groups using age and maximum oxygen consumption (VO_2_max) as stratifying factors (Fig. 4a). The age, weight, BMI, heart rate, and VO_2_max of each group are shown in the Supplementary Table 3. The three groups were fed with a placebo diet (placebo group), flaxseed lignans test food (FL group), or α-cyclodextrin test food (αCD group) for 9 weeks. Totally, 1, 2, and 2 subjects dropped out of the study for personal reasons in the placebo, FL, and αCD groups, respectively. Thus, analysis ultimately included data for 11, 10, and 10 subjects in the placebo, FL, and αCD groups. The rates of test food consumption in the placebo, FL, and αCD groups were 99.6 ± 1.0%, 97.1 ± 4.6%, and 97.8 ± 3.0%, respectively The test foods did not cause any adverse events during the experimental period.

**Fig. 4:**
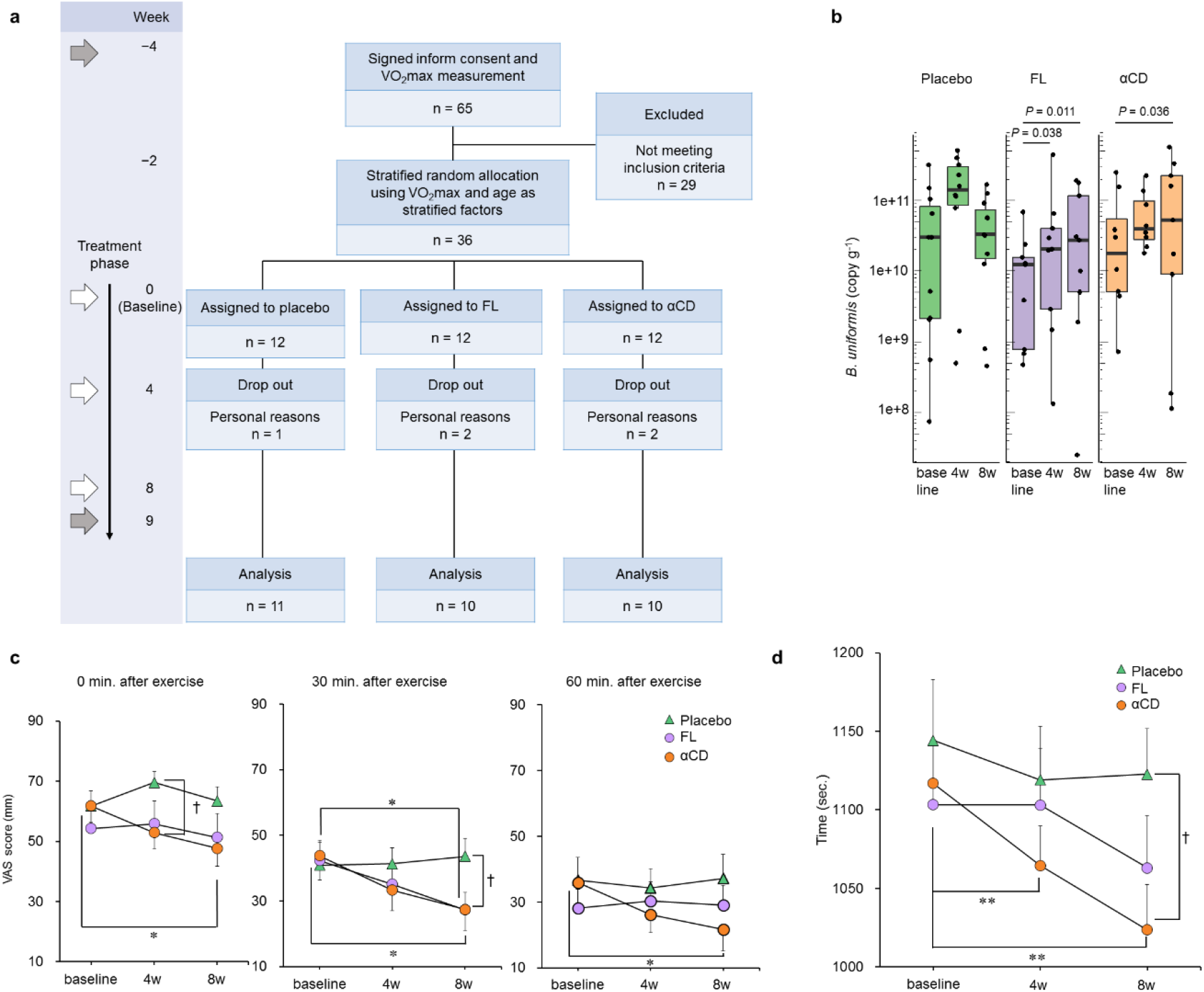
Flaxseed lignans (FL) and α-cyclodextrin (αCD) increased the cell number of *B. uniformis*, improved exercise performance and reduced post-exercise fatigue in human study. a. Human study protocol. VO_2_max was measured at the weeks indicated by the gray arrows. At the weeks indicated by the white arrows, we conducted a VAS questionnaire related to fatigue after 50 min of exercise, measured the time required to pedal 10 km on an exercise bike, and collected blood. b. Cell number of *B. uniformis* in feces were measured with qPCR at baseline and at 4 and 8 weeks of test food consumption. Data are presented as median with first and third quartiles as the box edges. The whiskers correspond to the minimum and maximum. Matched-pairs Wilcoxon signed-rank tests were performed to verify changes from baseline within each group. c. Results for a visual analog scale (VAS) questionnaire on fatigue after 50 min of exercise. Exercise tests were performed at 45% of each subject’s maximum exercise intensity. To determine statistical significance, unpaired or paired *t*-tests were used. d. Results for times required to pedal 10 km on an exercise bike. The intensity level (the weight of the pedal) was set at close to 45% of each subject’s maximum exercise intensity. Unpaired or paired *t*-tests were used to analysis. Values are expressed as mean ±s.e.m in panel c and d. All tests were two-tailed. †*P* < 0.05 (unpaired *t*-test). \**P* < 0.05 (paired *t*-test). \*\**P* < 0.01 (paired *t*-test).

At 0 (baseline), 4, and 8 weeks of test food consumption, we extracted DNA from the collected faecal samples, and performed quantitative PCR using *B. uniformis-specific* primers and probe. Neither the FL nor the αCD group exhibited a significant increase in *B. uniformis* cell number compared to that of the placebo group. However, in the FL group, the *B. uniformis* cell numbers were significantly higher at 4 and 8 weeks than at baseline; while in the αCD group, the cell number of *B. uniformis* was significantly higher at 8 weeks than at baseline (Fig. 4b). Despite these results, neither the FL nor the αCD group significantly differed with the placebo group in α or β diversity, suggesting that intake of flaxseed lignans or α-cyclodextrin did not cause any major changes in overall gut microbiota composition (Supplementary Fig. 2).

No significant difference in VO_2_max mean between before and after test food consumption (at 9 weeks) was noted in the placebo: 46.4 ± 1.9 versus 49.9 ± 2.9 (paired *t*-test; *P* = 0.118), FL group: 43.2 ± 2.2 versus 48.3 ± 3.8 (paired *t*-test; *P* = 0.060), or αCD group: 45.7 ± 2.7 versus 49.3 ± 3.9 (paired *t*-test; *P* = 0.186), respectively (Supplementary Table 4). Fig. 4c shows visual analogue scale (VAS) scores for fatigue following 50 min of constant-load bike-pedalling exercise. In the FL group, VAS scores at 0 min after exercise showed no significant change from baseline or no significant difference compared with the placebo group. However, at 30 min after exercise, VAS score in the FL group was significantly lower than that of baseline and the placebo group (8 weeks). In the αCD group, VAS scores after test food consumption (8 weeks) were significantly lower than baseline at 0, 30, and 60 min after exercise. Also, VAS score in the αCD group at 0 min after exercise at 4weeks consumption was significantly lower than the score obtained in the placebo group. Blood parameters measured in this study were shown in Supplementary Table 5. As for parameters in blood collected at immediately after exercise, the αCD group demonstrated a significant reduction in lactate levels after test food consumption (8 weeks: 16.53 ± 1.90 mg dl^-1^, mean ± s.e.m) compared to baseline (20.28 ± 2.30 mg dl^-1^, mean ± s.e.m, *P* = 0.007). Levels of diacron reactive oxygen metabolite (dROM), which represents oxidative stress, significantly increased in the placebo, FL, and αCD groups compared with baseline. We also measured blood concentrations of taurine, branched-chain amino acids (isoleucine, leucine, and valine), ornithine, citrulline and arginine since these amino acids may influence exercise performance and fatigue ^21–24^ However, there were no notable change in those amino acids. Also, no significant changes were observed in rate of perceived exertion (RPE), and muscle mass in the FL group or the αCD group, but there was slight increase of body fat mass in the αCD group (Supplementary Table 6-7). Fig. 4d shows the results of 10-km biking time-trial. The αCD group showed significantly shorter times at 4 weeks and after test food consumption (8 weeks) than at baseline, as well as a significantly shorter time than the placebo group after test food consumption (8 weeks).

## DISCUSSION

To our knowledge, this is the first study of dominant gut microbe *B. uniformis* in humans related to endurance exercise performance. In here, we showed improved endurance exercise ability in mice administered *B. uniformis* possibly due to enhancement of gluconeogenesis and β-oxidation in the liver through the high concentration of acetate and propionate in the cecum. We also noted that α-cyclodextrin increased the cell number of *B. uniformis* in the gut and improved exercise performance not only in mouse but also in human.

In our exercise tests, mice subjected to exercise after administering PBS exhibited a reduction in cecal SCFAs levels; whereas mice subjected to exercise following administration of *B. uniformis* did not exhibited the reduction. In addition, *B. uniformis*-monoassociated mice revealed that *B. uniformis* produced acetate and propionate in the intestine. The above findings suggested that the presence of *B. uniformis* in the gut resulted in an increase of acetate and propionate levels that exceeded those reduced by exercise, thereby maintaining their concentrations. Also, the liver of mice which exercised following *B. uniformis* administration showed elevated mRNA levels of *Cpt1a* and *Pck1*, which are rate limiting enzymes for β-oxidation and gluconeogenesis, respectively, indicating that β-oxidation and gluconeogenesis were activated in the murine liver. The expression of these genes has been reported to increase upon cecal infusion of propionate in the pig liver ^25^; meanwhile, acetate administration also increased expression of *Cpt1a* in the murine liver ^26^. Another study showed that over 60% of directly infused propionate in the cecum was used in glucose production, indicating it role as gluconeogenic substrate ^27^. Furthermore, NADH and ATP, produced by β oxidation of fatty acids in the liver, are used as energy sources for gluconeogenesis in the liver ^28,29^, suggesting that the acetate and propionate supplied by *B. uniformis* in the gut migrated to the liver and facilitated gluconeogenesis, resulting in the production of glucose needed for exercise. Especially, it was reported that intrarectal instillation of propionate improved endurance exercise performance in mice ^10^. Therefore, it is reasonable to consider the existence of a mechanism by which exercise performance is improved by the maintenance of propionate concentration in the gut as a result of propionate produced by *B. uniformis*. Meanwhile, according to Scheiman *et al*., lactate produced in the muscles due to exercise migrates to the gut where it is converted to propionate by *Veillonella atypica*^10^. However, no bacteria of that genus were detected in mice in our experiments. Thus, it is possible that the propionate observed in mice inoculated with *B. uniformis* in our study was not produced by bacteria of the genus *Veillonella* but by *B. uniformis*, using carbon sources from food. Few studies have examined changes in SCFAs concentration due to exercise, but a study reported that rats, which voluntarily exercised in a running wheel, contained similar concentrations of acetate and propionate to non-exercising rats; meanwhile, butyrate concentrations were higher in the rats subjected to exercise ^30^. In contrast to this reported study, our experiment observed that mice subjected to exercise following administration of PBS had a lower concentration of cecal SCFAs than mice not subjected to exercise, possibly due to a more intense than voluntary exercise and to immediate cecum collection after exercise. In a high-load exercise, SCFAs in the gut might be absorbed to obtain energy for exercise.

Bile acids are synthesized in hepatocytes conjugated with taurine or glycine and stored in the gallbladder. Conjugated bile acids secreted from the gallbladder into the duodenum are deconjugated and converted into secondary bile acids by gut bacteria ^31^, and these are reabsorbed in the terminal ileum and recirculated to the liver via the portal vein ^32^. While most of these bile acids are reused, some enter the bloodstream, migrate around the body, and function as signal molecules through the bile acid receptor TGR5 in multiple organs. TGR5 in skeletal muscle increases energy expenditure by promoting the conversion of inactive thyroxine into active thyroid hormone ^33^. In our experiments, the liver of mice administered with *B. uniformis*and got exercise contained higher levels of MCA, HDCA, and DCA than mice that were not subjected to exercise. However, there were no changes in total bile acid levels or in the expression of genes related to bile acid synthesis in the liver. While the function of the each bile acids as TGR5 agonists unclear, agonistic activity tends to be stronger with secondary bile acids than with primary bile acids ^34^. *B. uniformis* is able to hydrolyse many conjugated bile acids ^35^, therefore administration of the bacteria might promote changes in the composition of bile acids in the gut and bloodstream, resulting in changes in bile acid responses in muscles and thereby improving endurance.

We postulated two potential mechanisms by which *B. uniformis* improved endurance exercise performance (Fig. 5); however, further studies are required to determine the actual mechanism. Also, our experiments showed almost no changes in cecal concentrations of acetate or propionate, hepatic expression of *Cpt1a* or *Pck1*, or hepatic bile acid composition in *B. uniformis* mice or PBS mice without exercise. These findings suggested that these changes do not occur with *B. uniformis* administration alone and instead occur when exercise is added. In the future, we would investigate which type of signal (e.g., energy source depletion, increased blood flow, and stress) elicited the effects associated with *B. uniformis* inoculation.

**Fig 5:**
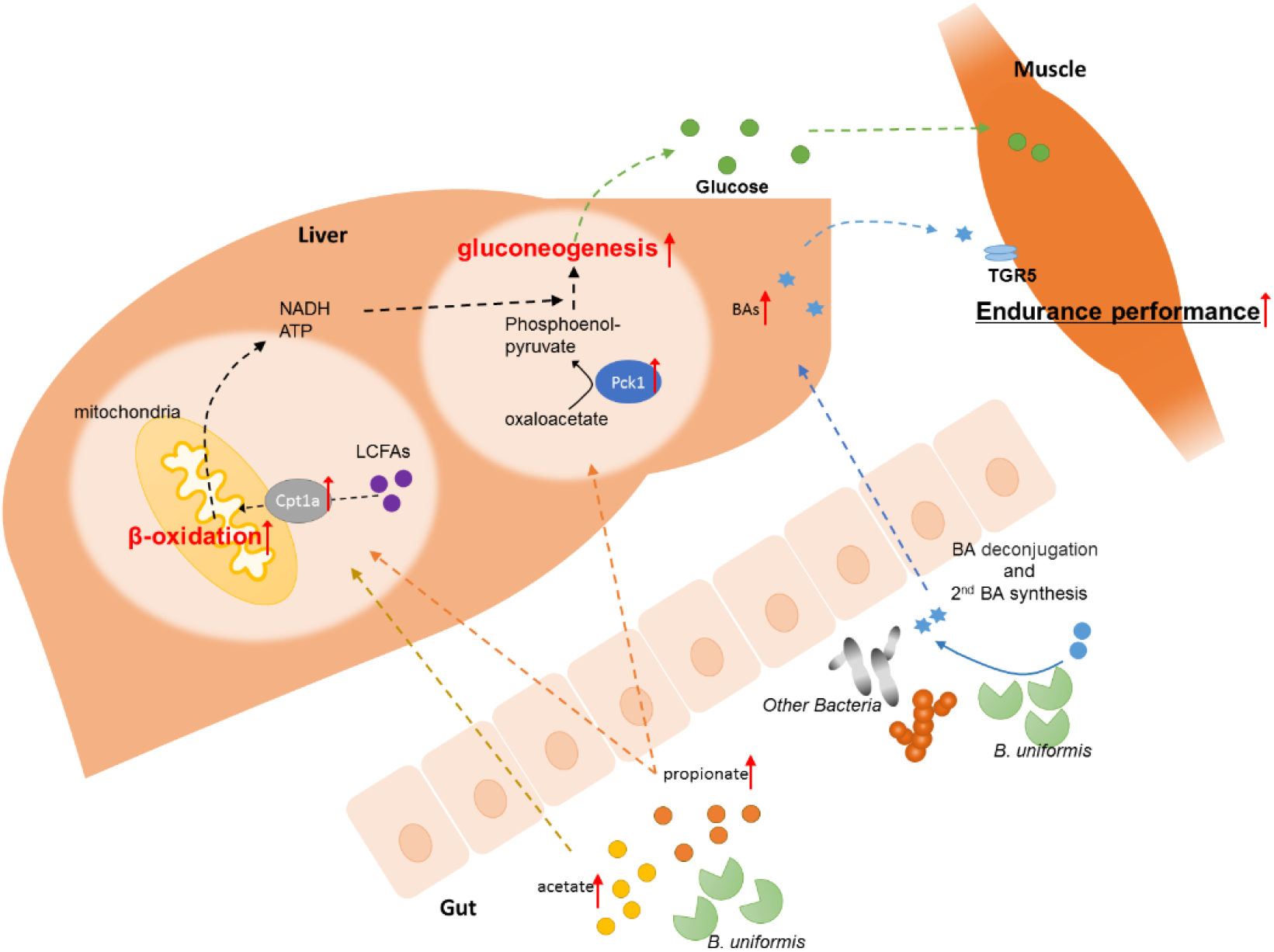
*B. uniformis* promotes host endurance performance through altering intestinal SCFAs production and BA composition. *B. uniformis* produces acetate and propionate in the intestine. These SCFAs migrate to the liver and increase expression of *Cptla* and *Pckl*. CPT1A transfers long chain fatty acids into mitochondria for β oxidation. PCK1 catalyzes the reversible conversion of oxaloacetate to phosphoenolpyruvate for gluconeogenesis. NADH and ATP produced by β oxidation of fatty acids in the liver are used as energy sources for gluconeogenesis in the liver. Acetate and propionate supplied by *B. uniformis* in the gut migrate to the liver and facilitate gluconeogenesis, resulting in the production of glucose needed for exercise. TGR5 in skeletal muscle increases energy expenditure by promoting the conversion of inactive thyroxine into active thyroid hormone. The liver of mice administered with *B. uniformis* subjected to exercise contained high levels of MCA, its secondary bile acid HDCA, and DCA, which may have promoted changes in the composition of bile acids in the bloodstream, resulting in changes in bile acid responses in muscles and thereby improving endurance. BA: bile acid; LCFAs: long chain fatty acids; TGR5; transmembrane G protein-coupled Receptor 5.

In this study, we also conducted a randomised, double-blind, placebo-controlled, parallel-group study to examine the effects of flaxseed lignans and α-cyclodextrin. Subjects who ingested α-cyclodextrin showed significantly elevated *B. uniformis* cell numbers in the gut compared with baseline, as well as reduced post-exercise fatigue and shorter times in time trials using exercise bike. These results suggested that α-cyclodextrin has a potential for enhance exercise performance through modification of gut microbiota in human. However, in our human study, as we did not administer *B. uniformis* cells directly to human due to the safety, we were unable to conclude that the same results obtained in the *B. uniformis* administered murine experiments also occurred in the human experiments even though α-cyclodextrin ingestion could enhance exercise performance and increase the abundance of *B. uniformis* in both human and mice.

In the present study, we determined that *B. uniformis*, a dominant bacterium in the human intestine, improved endurance exercise performance and we partially elucidated the causal relationship between *B. uniformis* and endurance exercise performance. The mechanisms by which *B. uniformis* improved endurance exercise performance relied on enhanced gluconeogenesis in the liver associated with acetate and propionate production and/or changes in bile acid composition. The findings of this study may contribute to a better understanding on how the gut microbiota relates to an increased endurance exercise performance and how the energy required for exercise is obtained.

## Materials and Methods

### Gut microbiome analysis of Japanese long-distance runners

Ethics statements: The study was approved by the Ethical Committee for Research in Humans of Shiba Palace clinic (Tokyo, Japan) in compliance with the Helsinki Declaration (Approval No. 2015AC-001). Forty-eight male athletes (age 18 to 22 years) from the Aoyama Gakuin University Track and Field Club Ekiden team, who lived in a dormitory, agreed to participate in the study in July 2015. Ten Japanese non-athlete males (age 22 to 24 years) also participated in the study, for comparison purposes. The protocol was approved by the Ethical Committee for Research in Humans of Aisei hospital Ueno clinic (Tokyo, Japan) in compliance with the Helsinki Declaration (Approval No. 27030). All the subjects agreed to participate in the study by providing informed consents prior to enrolment. If the participant was a minor, his parents provided consent.

Faecal microbiome analysis: Faecal samples from athletes and non-athletes were collected in test containers (Sarstedt K.K.) from July to September 2015 and September to October 2015, respectively The samples were kept at −30 °C for later use. Frozen samples were thawed at room temperature (approximately 24 °C), and then 0.2 g of faecal sample were aliquoted into plastic tubes. Samples were washed in 1.0 ml PBS buffer and then centrifuged at 14,000 × g for 3 min. Pellets were suspended in 500 μl extraction buffer (166 mM Tris/HCl, 66 mM EDTA, 8.3% SDS, pH 9.0) and 500 μl TE buffer-saturated phenol. Glass beads (300 mg, 0.1 mm diameter) were added to the suspension, and the mixture was vigorously vortexed for 60 sec using a Multi-beads Shocker (Yasui Kikai Corporation). After centrifugation at 14, 000 × g for 5 min, 400 μl of the supernatant were purified using phenol-chloroform-isoamyl alcohol, and DNA was precipitated with isopropanol. The samples were washed in 70% ethanol and dissolved in TE buffer. A High Pure PCR Template Preparation Kit (Roche Diagnostics K.K.) was used for further purification according to the manufacturer’s recommendations. Sequencing of the gene encoding 16S rRNA V4 region with MiSeq system and MiSeq Reagent Kit version 2 (300 Cycle) was carried out according to a previous report using 12.5 ng of DNA from each faecal sample ^36^. Five subjects in the athlete group and two subjects in the non-athlete group were excluded from analysis of gut microbiota because polymerase chain reaction for V4 region of bacterial 16S rRNA encoding gene did not yield sufficient amplification. QIIME was used for filtering and analysis of sequences ^37^. Quality filtering was performed using the provided fastq files, and sequences with a quality score under 29 were removed. Chimeric sequences were removed using USEARCH, and assignation of operational taxonomic units (OTUs) was carried out using open-reference OTU picking with a 97 % threshold of pairwise identity. Total number of sequence reads retained for analysis were 6,654,772. The mean, minimum, and maximum of the reads per sample were 130,485, 46,053, and 294,588, respectively. Alpha diversity (Chao1 richness estimates, Shannon diversity index, and phylogenetic distance [PD] whole tree) within two groups and the distances between subjects (unweighted and weighted UniFrac distances as a measurement of beta diversity) were also estimated using QIIME with rarefied data 35,000 reads per sample. Beta diversity was visualised by principal coordinate analysis. LEfSe analysis was performed in the following website: (http://huttenhower.sph.harvard.edu/galaxy/) ^38^. Quantitative real-time PCR was performed using species-specific primers and TaqMan probes, which are shown in Supplementary Table 8 and were previously reported ^39,40^. A LightCycler 480 Instrument II (Roche Diagnostics K.K.) was used for the amplification and detection of specific sequences. The standard curve was created by amplification of 10-fold serial dilutions of bacterial 16S rRNA sequence, synthesised by Thermo Fisher Scientific K.K.

Blood analyses: Blood samples from athletes and non-athlete men were collected during the same period as the faecal samples. Different tubes were used based on the analyses of interest. Ethylenediaminetetraacetic acid disodium salt dihydrate-coated tubes for Hb, MCH, MCHC, MCV, hematocrit, and RBC; non-coated tubes for CPK, Fe and ferritin. The collected blood samples were sent directly to the clinical laboratory.

A 3,000-m race: Measurement of the time taken to run 3,000 m in the field track of Aoyama Gakuin University was recorded. Twenty-five out of 48 athletes participated in this trial.

### Animals and sample collection

*B. uniformis* culture: *B. uniformis* JCM5828^T^ was cultured on BL agar plate (Nissui Pharmaceutical Co., Ltd.) for 48 h at 37 °C under anaerobic condition. Colonies obtained from BL agar plates were collected and incubated overnight in 25 ml of GAM broth (Nissui Pharmaceutical) at 37 °C under anaerobic condition. To measure the number of viable bacteria cells, bacterial culture was diluted as appropriate and plated onto BL agar plates. After 48 h culture, the numbers of colonies formed on the plates were counted, and colony forming unit (CFU) ml^-1^ was calculated.

Exhaustive Swimming Test: Specific-pathogen-free male C57BL/6J mice (Japan SLC, Inc.) were housed singly, and maintained on a 12 h light/dark cycle at room temperature (22±3°C) with food and water provided *ad libitum*. Mice were fed with a control diet CE-2 (Oriental Yeast Co., Ltd.), (Fig. 2) or CE-2 containing 5% (w w^-1^) of test substance (flaxseed lignans or α-cyclodextrin) (Fig. 3). Nippn flaxseed lignans (Nippon Flour Mills Co., Ltd.) and Dexypearl-α (Ensuiko Sugar Refining Co., Ltd.) were used as flaxseed lignans and α-cyclodextrin, respectively. All experimental groups were orally administered with *B. uniformis* JCM5828^T^ (2×10^8^ CFU, 200 μl of PBS per mouse) for 6-10 consecutive weeks, while the PBS groups were treated with same volume of sterile PBS. The faecal samples were collected at day 0 and at the end of the experiment and immediately stored at −80 °C. Faecal DNAs were prepared from 50 mg of faecal samples by using the same protocol for human faecal sample described above. All mice were sacrificed under isoflurane anaesthesia after last swimming endurance test or swimming training, and whole blood was collected from the inferior vena. The liver and cecal contents were frozen immediately in liquid nitrogen and stored at −80 °C. All experiments were performed following approval by the Institutional Animal Care and Use Committee of Sankyo Labo Service Corporation, Inc, New Drug Research Center Inc, or Asahi quality and innovations, Ltd. (Fig. 2c, d, f, g: Approval No.; SKL-A-17095, Approval date; 22nd February, 2018, Experimental period; From February to May, 2018. Fig. 2a, b: Approval No.; 170428A, Approval date; 1st May, 2017, Experimental period; From May to July, 2017. Fig. 3a-d: Approval No.; 171010D, Approval date; 17th October, 2017, Experimental period; From October to December, 2017. Fig. 3e-h: Approval No.; 18-21-01, Approval date; 11th May, 2018, Experimental period; From May to August, 2018).

*B. uniformis*-monoassociated mice: Germ-free grade male C57BL/6N mice (Sankyo Labo Service Corporation, Inc) were housed 3-4 per cage and maintained in a vinyl isolator in a room kept at a constant temperature (23±2°C). The *B. uniformis* GB group (BU GB) was treated with a single dose of *B. uniformis* JCM5828^T^ (2×10^9^ CFU, 200 μl of PBS per mice), and control group (GF) was treated with same volume of sterile PBS. After 2 weeks, all mice were sacrificed under isofurane anaesthesia. Fecal samples were collected at day0 and last day of experiment and immediately stored at −80 °C. The experiment was performed following approval by the Institutional Animal Care and Use Committee of Sankyo Labo Service Corporation, Inc (Fig. 2e: Approval No.; SKL-J18004, Approval date; 20th April, 2018, Experimental period; From April to May, 2018).

Swimming test (Exercise protocol): In order to evaluate the physical performance, mice were exercised in a pool developed at Kyoto University (ANITEC CO., LTD) ^41,42^. Mice had an acclimation swimming period 3 weeks before the exhaustive swimming test. The acclimation swimming was performed every 3 days and consisted of the following: Day 1 swimming 8 l min^-1^ flow rate, increased by 1 l min^-1^ every 3 minutes, and ended by swimming at 12 l min^-1^ for 3 minutes; Day 2 swimming 10 l min^-1^ for 5 min, increased by 1 l min^-1^every 5 minutes, and finished after swimming at 12 l min^-1^ for 5 min; and Day 3 swimming 12 l min^-1^ for 5 min, followed by 13 l min^-1^ for 15 minutes, and finished after swimming for 14 l min^-1^ for 15 minutes. Two or 3 days after the swimming practice, swimming endurance capacity was evaluated. It started at 12 l min^-1^ for 5 min, followed by 13 l min^-1^ for 5 min, and finished with exhaustion at 14 l min^-1^. The swimming endurance time of each mouse was recorded from beginning to exhaustion, which was determined by failure to return to the surface within 3 seconds. The grouping was performed by stratified continuous randomisation, or so that the average swimming time of each group was equal. During the experimental term, mice were subjected to weekly swimming test. Endurance swimming test were performed at the same condition as grouping (Fig. 2a,b, Fig. 3). During training, mice were exercised for 15 min (Swimming 12 l min^-1^ for 5 min, followed by 13 l min^-1^ for 10 min. Fig 2. c-d, f, g).

RNA sequencing: Six samples were selected randomly from each group. Total RNA from liver was prepared using RNeasy Kit (Qiagen), and the subsequent steps were performed by Bioengineering Lab. Co., Ltd. For cDNA library construction, KAPA Stranded mRNA-Seq Kit (KAPA BIOSYSTEMS) and Fast Gene Adapter Kit (Fast Gene) were used, according to the manufacturer’s protocol. The library fragments were sequenced on the HiSeqX (Illumina) system. For data analysis, read sequences (75 bp) were aligned to the Mus musculus (GRCm38.p6) reference genome (https://www.ncbi.nlm.nih.gov/assembly/GCF_000001635.26/) using hisat2. To identify genes differentially expressed, the iDEGES/edgeR were used. GO analysis was performed with DAVID website (https://david.ncifcrf.gov/)^43^.

Reverse transcriptional quantitative PCR (RT-qPCR): Eight samples were selected randomly for RT-qPCR from each group. Total RNA was prepared using RNeasy Kit (Qiagen) according to the instructions of the manufacturer. cDNA was synthesized using PrimeScript RT reagent Kit (Takara Bio Inc.) RT-qPCR was performed using LightCycler 480 system (Roche Diagnostics K.K.) and TB Green Premix Ex Taq II (Takara Bio Inc.). The sequences of *Pck1* and *Cpt1a* primers were referred to a previous report ^44,45^. The relative amounts of these mRNAs were normalized using the *Actb* mRNA signal.

SCFAs analysis: Measurements of the SCFAs were performed by Techno Suruga Laboratory Co., Ltd. One hundred mg of samples were pre-treated according to previously reported method ^46^. After pretreatment, SCFAs levels were measured by Gas Chromatography-Flame Ionisation Detector (GC-FID) system (7890B, Agilent Technologies) with DB-WAXetr column (30 m, 0.25 mm id, 0.25 μm film thickness).

Bile acids analysis: Bile acid levels were measured as previously described with some modifications ^47^. Specifically, 20 mg of liver tissue were mixed with 300 μL of 50% methanol in water (v v^-1^) containing an internal standard (20 μM CSA), homogenised with a multi-sample homogeniser (Shake Master Neo, Bio Medical Science), and centrifuged. The samples were centrifuged at 20,400 ×*g* for 10 min at 10 °C, and supernatants were collected. Liquid chromatography separation was performed using an Agilent 1290 UPLC system (Agilent Technologies), an ACQUTTY UPLC HSS T3 column (1.8 μm, 50 mm × 2.1 mm ID; Waters), and a mobile phase flow rate of 0.3 ml min^-1^ at 45 °C. Gradient elution mobile phases consisted of A (water, 50 mM ammonium formate) and B (methanol, 50 mM ammonium formate). The initial gradient consisted of 10% B, and linearly increased to 50% for 0.1 min, then maintained at 50% B for 2 min, linearly increased again to 80% for 9.5 min, and then linearly increased to 100% for 10.5 min; maintained for 12 min, with subsequent reequilibration with 10% B for a 5 more min. Sample temperature was maintained at 4 °C in the autosampler prior to analysis. The sample injection volume was 1 μl. Mass spectrometer (MS) analysis was performed using an Agilent 6490 triple quadrupole mass spectrometer MS (Agilent Technologies) equipped with an Agilent Jet Stream ESI probe in negative-ion mode. A capillary voltage of −3,500 V, a source temperature of200 °C, gas flow of 14 l min^-1^, nebuliser of 50 psi, sheath gas temperature of250 °C, and sheath gas flow of 11 l min^-1^ were used. Concentrations of individual bile acids were calculated from the peak areas in the chromatogram detected with MRM relative to the internal standard, CSA.

### Human study with flaxseed lignan or α-cyclodextrin as test foods

Ethic statement: This study was conducted in accordance with the Declaration of Helsinki and Ethical Guidelines for Medical and Health Research Involving Human Subjects (notification of the Ministry of Education, Culture, Sports, Science and Technology, and the Ministry of Health, Labour and Welfare). The study protocol was reviewed prior to implementation by the Institutional Review Board of Chiyoda Paramedical Care Clinic and was implemented upon approval (Approval No: 18071903, approval date: 20th July, 2018). The study was also registered in the University Hospital Medical Information Network system (Registration ID: UMIN000033748, registration date: 14th August, 2018). Informed consent was provided by all participants prior to preliminary test which started on 17th August, 2018..

Subjects: Thirty-six Japanese men (12 per group), who met all the following inclusion but not exclusion criteria, consented to participate in writing, based on free will, after receiving a thorough explanation of the nature of the experiment. This study was conducted as a pilot study; therefore, no statistical evidence for the sample size was available. Inclusion criteria: (1) healthy men aged 20-49 years at the time of consent; (2) having exercise habits. The 1-2 sessions (≥ 30 min per session) per week of exercise of ≥ 5 metabolic equivalents (METs), excluding resistance (weight) training, among the exercises listed in the 2011 Compendium of Physical Activities: A Second Update of Codes and MET Values published by the National Institute of Health and Nutrition; (3) continuation of exercise habits during the study period; and (4) able to refrain from consuming prohibited foods for at least 1 week prior to the date of consent or at any point during the study period. Exclusion criteria: (1) self-report of ≤ 5 bowel movements per week, having a “weak stomach”, or being prone to diarrhoea; (2) past or current serious heart, liver, kidney, or gastrointestinal disease; (3) high alcohol consumption; (4) extremely irregular eating habits or irregular daily rhythm due to shift or night work; and (5) deemed unsuited to participate in the experiment by the principal investigator or a sub-investigator.

Study Design: The experiment was conducted as a randomised, double-blinded, placebo-controlled, parallel-group study From 5 weeks to 3 weeks prior to intake of test foods, VO_2_max was measured in 65 individuals whose participation in the study was assured in advance and who consented to participate. With VO_2_max and age as stratifying factors, subjects were randomly assigned into three groups (12 subjects/group) to minimise bias. All subjects ingested three tablets of test food daily for 9 weeks from the first day of the experiment. Subjects were subjected to designated tests at baseline (0) and at 4, 8, and 9 weeks at the Chiyoda Paramedical Care Clinic. Subjects received prescribed meals for dinner on days prior to visiting the clinic and for breakfast on the days of clinic visits. Gut flora, free fatty acids, and dROM were measured by Asahi quality and innovations, Ltd., while all other measurements and tests were performed by Chiyoda Paramedical Care Clinic Co. Ltd. (Tokyo, Japan). Safety was monitored and adverse events were recorded during the assessment of hematology and clinical chemistry results, and vital signs of each participant, in addition to physical examinations at each clinic visit and. This study was conducted from 20th July to 25th December, 2018.

Test foods: The test food was Nippn flaxseed lignans (Nippon Flour Mills Co., Ltd.), which was chosen as a useable form of SDG. Nippn flaxseed lignans contained approximately 40% (w w^-1^) SDG and 40% (w w^-1^) α-cyclodextrin. Tablets containing Nippn flaxseed lignans were designated as FL test food. The α-cyclodextrin test food contained Dexipearl-α (Ensuiko Sugar Refining Co., Ltd.), designated αCD. The placebo tablet did not contain neither flaxseed lignans nor α-cyclodextrin. Supplementary Table 9 shows the compositions of FL test food, αCD test food, and the placebo.

Testing Methods: 1) VO_2_max and ventilatory threshold (VT); exercise testing was conducted with a stationary bicycle (AEROBIKE-75XLIII; Konami Sports Life). Gas expired during exercise was measured with an Aero Monitor expired gas analyser (AE-310s; Minato Medical Science Co., Ltd.). The exercise load was incrementally increased by 15 W every minute until exhaustion. Concentrations of oxygen and carbon dioxide in expired gas were measured with the Aero Monitor. During exercise, subjects maintained a pedal speed of 60 rpm. Heart rate was measured with a monitor during exercise. Exhaustion was determined based on the following criteria: i) the subject declared himself to be exhausted or was unable to continue exercising; ii) the subject’s heart rate reached the estimated maximum heart rate for his age, defined as 220 - age; or iii) the subject was declared exhausted according to a physician evaluation. The maximum oxygen consumption and the weight of the pedal immediately before fulfilling one of the above three criteria were defined as VO_2_max and maximum exercise intensity, respectively. 2) Fatigue questionnaire and RPE; Exercise testing was carried out with a stationary bicycle as described above. Subjects were subjected to exercise at 45% of their maximum exercise intensity for 50 min. During exercise, subjects maintained a pedal speed of 60 rpm. RPE was measured with the Borg scale at baseline and at 10, 20, 30, 40, 45, and 50 min of exercise. In addition, a VAS questionnaire was used to assess fatigue after exercise testing (0, 30, 60 min). The VAS questionnaire used in this study was created by the Japanese Society of Fatigue Science (http://www.hirougakkai.com/). 3) Exercise performance; Measured by the time required to pedal 10 km on a stationary bicycle (FBS-101; Fujimori Ltd.). The intensity level (i.e., the weight of the pedal) was set close to 45% of maximum exercise intensity. 4) Blood testing (lactate, CPK, lactate dehydrogenase (LDH), glucose (GLC), creatine, free fatty acids, cortisol, amino acids, glucagon, IL6, growth hormone, and dROM); different tubes were used based on the analyses of interest. Ethylenediaminetetraacetic acid disodium salt dihydrate-coated tubes for amino acids; aprotìnin-coated tubes for glucagon; heparin-coated tube for GLC; perchloric acid-added tube for lactate; non-coated tubes for CPK, creatine, LDH, free fatty acids, cortisol, IL6, growth hormone, and dROM. Blood was collected immediately before, after, and 60 min after exercise to assess fatigue and RPE. Blood collected immediately before and after exercise testing was used to measure lactate, CPK, LDH, GLC, creatine, free fatty acids, cortisol, amino acids, glucagon, IL6, growth hormone, and dROM. Blood collected 60 min after exercise testing was used to measure only LDH, CPK, creatine, GLC, and lactate. 5) Muscle mass and body fat mass; it was measured with a body composition analyser (InBody 570; InBody Inc.) immediately before exercise to assess fatigue and RPE. 6) Gut flora; Stool samples were collected at baseline and after 4 and 8 weeks of test food consumption. The stool samples were immediately frozen and preserved at −30 °C until use. As described above, DNA was extracted from stool samples, and gut flora were analysed with MiSeq. Total number of sequence reads retained for analysis were 4,757,361. The mean, minimum, and maximum of the reads per sample were 52,859, 12,805, and 125,205, respectively. To estimate alpha and beta diversity, rarefied data of 10,000 reads per sample were used. Cell number of *B. uniformis* was determined using specie-specific qPCR, as described above. One and two subjects were excluded from FL and αCD group, respectively, in the comparison of the cell number of *B. uniformis* because the cell numbers were under detectable level.

### Statistical analyses

In the athlete study, to compare the gut microbial alpha and beta diversities between groups, we used a two-sample *t*-test using Monte Carlo permutations and the permutational multivariate analysis of variance (PREMANOVA), respectively. The number of permutations for both tests was set to 999 to calculate *P*-values. Correlation between the relative abundance of *Bacteroides*, *Lachnospira*, *Sutterella*, *Escherichia*, *Gemellaceae*, *Streptophyta* in the faecal samples from the athletes and, blood data or 3,000-m race time of the athletes were determined using the Spearman rank correlation coefficient. Correlation between the log10 cell numbers of each *Bacteroides*-species in the fecal sample from the athletes and the 3,000-m race time were determined using the Pearson correlation coefficient. Mann-Whitney *U*-tests were used to investigate differences in the relative abundance of genera and log10 cell number of each *Bacteroides-species* between athlete group and non-athlete group. In mice study, differences of swimming time to exhaustion between groups was assessed by using the Mann-Whitney *U*-test. SCFAs in cecum or transcriptional levels of *Cpt1a* and *Pck1* were tested by Tukey-Kramer. SCFAs in feces were tested by unpaired *t*-test. False discovery rates (*q*-value) of RNA sequencing data were estimated by using the Benjamin-Hochberg procedure. In the human study using FL or αCD as test foods, Mann-Whitney *U*-tests assessed the differences of *B. uniformis* cell numbers between the two treated groups versus the placebo group, and Wilcoxon signed-rank tests were performed to identify the differences between baseline and 4 or 8 weeks within each group. To determine statistical significance for other parameters in the two treated groups versus the placebo group, or the baseline and 4 or 8 weeks within groups, unpaired or paired *t*-tests were used, respectively Data were analysed with the SPSS (ver. 23, IBM Japan, Ltd.) or R (ver. 3.4.2 and 3.6.0). Differences with *P* < 0.05 or 0.01 were considered statistically significant. Data are expressed as mean±s.e.m., mean±s.d., or median±s.e.m.

## Acknowledgements

We thank Kazuto Washida and Yasuhisa Miyamoto (Asahi quality and innovations Co., Ltd.) for measuring alpha cyclodextrin in the Nippn amanilignan. We also thank Kyoko Morimoto for her excellent technical assistance. This study was supported in part by JSPS KAKENHI (18H04805 to S.F.), JST PRESTO (JPMJPR1537 to S.F.), JST ERATO (JPMJER1902 to S.F.), AMED-CREST (JP19gm1010009 to S.F.), the Takeda Science Foundation (to S.F.), the Food Science Institute Foundation (to S.F.), and the Program for the Advancement of Research in Core Projects under Keio University’s Longevity Initiative (to S.F.).

## Author contributions

H.M., C.K., T.N., and S.F. designed the study. S.F. and T.N. managed the whole project. C.K. and H.M. performed animal and clinical experiments. S.H. and Y.C. supervised the athlete study. C.I., N.K., and T.I. conducted metabolome analyses. H.M., C.K., and S.F. wrote the manuscript, and all authors reviewed the manuscript.

