## supplemental figures and tables for "*Bacteroides uniformis* enhances endurance exercise performance through gluconeogenesis"

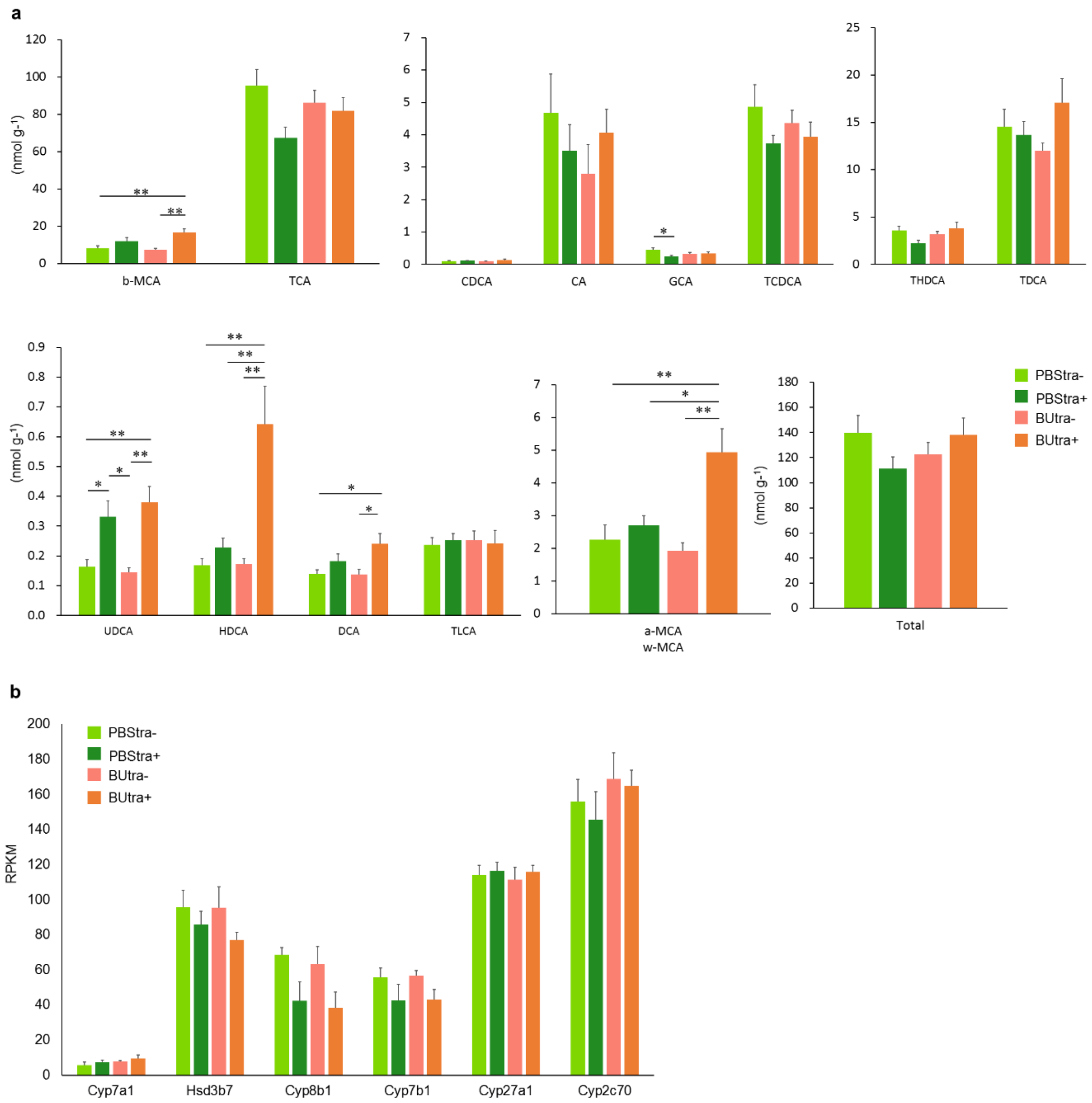


Supplementary Fig1. Hepatic bile acid composition and expression of genes related to bile acid production.

**a**, Hepatic bile acid composition. β-MCA, UDCA, HDCA, DCA, and α-MCA and ω-MCA levels were significantly higher in BUtra+ than in BUtra-. HDCA, and α-MCA and ω-MCA levels were also significantly higher than in PBStra+. GCA levels were significantly low in PBStra+. UDCA levels were significantly higher in PBStra+ and BUtra+ than in the respective groups of mice that were not subjected to exercise. **b**, Comparison of expression of genes related to bile acid production. No significant differences in expression were observed for any genes among the four groups. Data are shown as mean ± s.e.m. Statistical significance of difference between groups were analysed by Tukey-Kramer test. ** *P*<0.01 , **P*<0.05

PBStra+: mice administered PBS and subjected to exercise; PBStra-: mice administered PBS and not subjected to exercise; BUtra+: mice administered *B. uniformis* and subjected to exercise; BUtra-:

mice administered *B. uniformis* and not subjected to exercise. α-MCA: α-muricholic acid; β-MCA: β-muricholic acid; HDCA: hyodeoxycholic acid; DCA: deoxycholic acid; CA: cholic acid; TCA: taurocholic acid; CDCA: chenodeoxycholic acid; TCDCA: taurochenodeoxycholic acid; GCA: glycocholic acid; THDCA: taurohyodeoxycholic acid; TDCA: taurodeoxycholic acid; UDCA: ursodeoxycholic acid; TLCA: taurolithocholic acid.

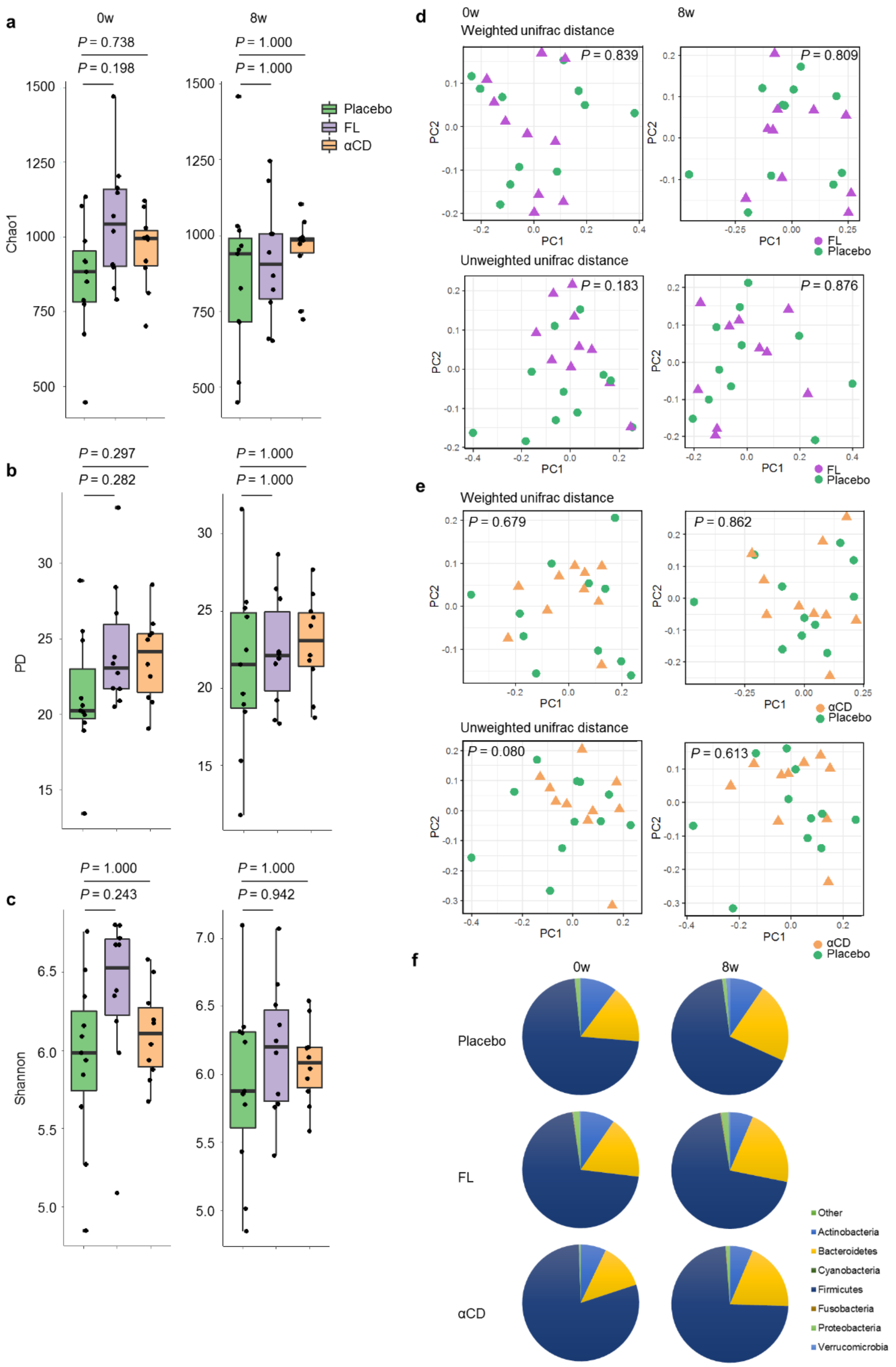


Supplementary Fig. 2: Supplementation of flaxseed lignan or α-cyclodextrin does not lead to overall gut microbiome alteration in human.

**a**-**c**, Comparison of Chao1(**a**), PD whole tree (**b**), and Shannon diversity index (**c**) between placebo and treated groups. Data are presented as median with first and third quartiles as the box edges. The whiskers correspond to the minimum and maximum. Statistical significance of differences between groups was analyzed by a two-sample *t*-test using Monte Carlo permutations. **d-e**, Comparison of microbiota β diversity between placebo and FL group (**d**) and αCD group (**e**) on weighted and unweighted UniFrac distance. Statistical significance of differences between groups was analysed by the PREMANOVA. **f**, Gut microbial composition of placebo and treated groups. Mean relative abundance of each bacterial phyla are shown in pie charts. Placebo, $n$ = 11; FL group $n$ = 10; αCD group $n$ = 10. PREMANOVA: permutational multivariate analysis of variance.

Supplementary Table 1. Relative abundance of 9 genera picked up by LEfSe analysis in athlete and non-athlete.

| Genus | Mean ± s.e.m (%) | | | | | | Median ± s.e.m (%) | | | | | | *P*-value | Detection rate(%) [1)] | |
|---|---|---|---|---|---|---|---|---|---|---|---|---|---|---|---|
| | Athlete | | | Non-athlete | | | Athlete | | | Non-athlete | | | | Athlete | Non-athlete |
| *Bacteroides* | 12.63 | ± | 1.68 | 8.49 | ± | 4.55 | 11.41 | ± | 1.68 | 1.23 | ± | 4.55 | 0.014 | 100.0 | 100.0 |
| *Prevotella* | 1.01 | ± | 0.34 | 0 | | | 0 | ± | 0.34 | 0 | | | 0.067 | 37.2 | 0.0 |
| *Lachnospira* | 0.56 | ± | 0.10 | 0.26 | ± | 0.14 | 0.28 | ± | 0.10 | 0.025 | ± | 0.14 | 0.022 | 93.0 | 50.0 |
| *Sutterella* | 0.41 | ± | 0.13 | 0.30 | ± | 0.17 | 0.10 | ± | 0.13 | 0.010 | ± | 0.17 | 0.021 | 90.7 | 62.5 |
| *Escherichia* | 0.37 | ± | 0.18 | 0.67 | ± | 0.32 | 0.010 | ± | 0.18 | 0.32 | ± | 0.32 | 0.002 | 60.5 | 100.0 |
| *Granulicatella* | 0.0012 | ± | 0.0005 | 0.0047 | ± | 0.0018 | 0 | ± | 0.0005 | 0 | ± | 0.0018 | 0.099 | 11.6 | 37.5 |
| Gemellaceae;g_ | 0.00093 | ± | 0.00045 | 0.0056 | ± | 0.0025 | 0 | ± | 0.00045 | 0.010 | ± | 0.0025 | 0.001 | 9.3 | 62.5 |
| Streptophyta;f_;g_ | 0.00023 | ± | 0.00023 | 0.011 | ± | 0.006 | 0 | ± | 0.00023 | 0 | ± | 0.006 | 0.008 | 2.3 | 37.5 |
| *Citrullus* | 0 | | | 0.0022 | ± | 0.0013 | 0 | | | 0 | ± | 0.0013 | 0.157 | 0.0 | 12.5 |

Comparison of the relative abundance between groups were conducted with Mann–Whitney *U*-tests.

Supplementary Table 2. Results of species-specific qPCR using DNAs extracted from feces of athlete and non-athlete.

| | Median log10 cell number ± s.e.m. (No. of 16S rRNA gene/g) | | | No. of detected samples/ total no. of samples (%) | | Correlation with 3,000-m race time | |
|---|---|---|---|---|---|---|---|
| | Athlete | Non-athlete | *P*-value | Athlete | Non-athlete | Correlation coefficient | *P*-value |
| *Bacteroides caccae* | 6.81 ± 0.17 | 6.05 ± 0.26 | 0.036 | 100.0% | 100.0% | 0.06 | 0.760 |
| *Bacteroides eggerthii* | 7.04 ± 0.15 | 6.24 ± 0.11 | 0.005 | 91.7% | 100.0% | 0.03 | 0.897 |
| *Bacteroides uniformis* | 10.52 ± 0.15 | 9.38 ± 0.20 | 0.001 | 89.6% | 100.0% | -0.53 | 0.011 |
| *Bacteroides thetaiotaomicron* | 9.49 ± 0.13 | 8.05 ± 0.30 | 0.006 | 89.6% | 90.0% | -0.24 | 0.280 |
| *Bacteroides vulgatus* | 10.61 ± 0.19 | 9.74 ± 0.23 | 0.037 | 85.4% | 90.0% | 0.00 | 0.990 |
| *Bacteroides dorei* | 10.17 ± 0.14 | 9.21 ± 0.22 | 0.009 | 75.0% | 60.0% | -0.09 | 0.698 |
| *Bacteroides fragilis* | 9.67 ± 0.15 | 8.95 ± 0.36 | 0.651 | 50.0% | 30.0% | N.T. | N.T. |
| *Bacteroides stercoris* | 9.94 ± 0.24 | 9.04 ± 0.43 | 0.168 | 47.9% | 40.0% | N.T. | N.T. |
| *Bacteroides plebeius* | 10.20 ± 0.17 | 9.29 | N.T. | 29.2% | 10.0% | N.T. | N.T. |
| *Bacteroides intestinalis* | 8.63 ± 0.35 | 8.80 | N.T. | 20.8% | 10.0% | N.T. | N.T. |
| *Bacteroides finegoldii* | 9.64 ± 0.42 | N.D. | | 12.5% | 0.0% | N.T. | N.T. |
| *Bacteroides coprophilus* | 10.00 ± 0.23 | N.D. | | 12.5% | 0.0% | N.T. | N.T. |
| *Bacteroides coprocola* | 4.14 ± 0.59 | N.D. | | 4.2% | 0.0% | N.T. | N.T. |

N.D. : not detectable, N.T. : not tested

Comparison of the cell number between groups were conducted with Mann–Whitney *U*-tests.

Correlation between bacterial cell numbers and 3,000-m race time in athlete group were assessed with the Pearson correlation coefficients.

Supplementary Table 3. Subjects background in human study.

| | Placebo (*n* = 11) | FL (*n* = 10) | αCD (*n* = 10) | *P*-value 1) | |
|---|---|---|---|---|---|
| | | | | Placebo vs. FL | Placebo vs. αCD |
| Age | 36.3 ± 9.6 | 33.9 ± 10.0 | 34.5 ± 10.9 | 0.59 | 0.70 |
| Weight (kg) | 64.01 ± 9.68 | 63.12 ± 6.34 | 67.76 ± 7.54 | 0.81 | 0.34 |
| Body mass index (kg/$m^2$) | 22.01 ± 2.43 | 21.53 ± 1.67 | 22.23 ± 2.43 | 0.61 | 0.84 |
| Heart rate (bpm) | 72.5 ± 9.8 | 70.5 ± 13.0 | 69.1 ± 9.4 | 0.70 | 0.43 |
| $VO_2$max (ml/kg/min) | 46.40 ± 6.45 | 43.20 ± 6.83 | 45.67 ± 8.66 | 0.28 | 0.83 |

Mean ± standard deviation

1)two-tailed unpaired *t*-test

Supplementary Table 4. $VO_2max$ and ventilator threshold (VT) at baseline and after consumption (9 weeks).

| | Group | Baseline | 9 weeks | *P*-value (vs. baseline)[1)] | *P*-value (vs. placebo)[2)] | |
|---|---|---|---|---|---|---|
| | | | | | Baseline | 9 weeks |
| $VO_2max$ (ml/kg/min) | Placebo | 46.40 ± 1.94 | 49.93 ± 2.88 | 0.12 | - | - |
| | FL | 43.20 ± 2.16 | 48.31 ± 3.77 | 0.06 | 0.28 | 0.73 |
| | αCD | 45.67 ± 2.74 | 49.30 ± 3.90 | 0.19 | 0.83 | 0.90 |
| VT (ml/min) | Placebo | 857.0 ± 56.3 | 874.2 ± 56.7 | 0.66 | - | - |
| | FL | 811.4 ± 49.6 | 852.7 ± 66.7 | 0.46 | 0.55 | 0.81 |
| | αCD | 939.7 ± 44.3 | 932.1 ± 54.9 | 0.84 | 0.27 | 0.47 |

Mean ± standard error

1) two-tailed paired *t*-test

2) two-tailed unpaired *t*-test

Supplementary Table 5. Hematological parameters at baseline, 4 and 8weeks in human study.

| | Unit | Group | Time point | Baseline | 4 weeks | 8 weeks | *P*-value (baseline vs.) [1)] | | *P*-value (placebo vs.) [2)] | | |
|---|---|---|---|---|---|---|---|---|---|---|---|
| | | | | | | | 4 weeks | 8 weeks | Baseline | 4 weeks | 8 weeks |
| Creatine | mg/dl | Placebo | before exercise | 0.30 ± 0.07 | 0.33 ± 0.06 | 0.32 ± 0.04 | 0.39 | 0.71 | - | - | - |
| | | | after exercise | 0.45 ± 0.10 | 0.50 ± 0.10 | 0.47 ± 0.07 | 0.18 | 0.74 | - | - | - |
| | | | 60 min after exercise | 0.31 ± 0.08 | 0.34 ± 0.08 | 0.30 ± 0.05 | 0.34 | 0.85 | - | - | - |
| | | FL | before exercise | 0.31 ± 0.05 | 0.24 ± 0.05 | 0.32 ± 0.06 | 0.11 | 0.80 | 0.91 | 0.31 | 0.98 |
| | | | after exercise | 0.47 ± 0.06 | 0.45 ± 0.07 | 0.50 ± 0.08 | 0.59 | 0.43 | 0.90 | 0.68 | 0.79 |
| | | | 60 min after exercise | 0.32 ± 0.04 | 0.25 ± 0.05 | 0.33 ± 0.06 | **0.02** | 0.76 | 0.91 | 0.36 | 0.70 |
| | | αCD | before exercise | 0.39 ± 0.11 | 0.43 ± 0.11 | 0.41 ± 0.09 | 0.34 | 0.74 | 0.49 | 0.42 | 0.35 |
| | | | after exercise | 0.56 ± 0.14 | 0.59 ± 0.13 | 0.55 ± 0.11 | 0.62 | 0.90 | 0.54 | 0.58 | 0.55 |
| | | | 60 min after exercise | 0.40 ± 0.11 | 0.45 ± 0.11 | 0.42 ± 0.09 | 0.32 | 0.74 | 0.52 | 0.41 | 0.26 |
| Creatine phosphokinase | U/l | Placebo | before exercise | 141.82 ± 20.24 | 122.7 ± 14.08 | 141.4 ± 21.91 | **0.03** | 0.97 | - | - | - |
| | | | after exercise | 158.18 ± 22.64 | 133.3 ± 14.63 | 151.3 ± 22.94 | **0.03** | 0.66 | - | - | - |
| | | | 60 min after exercise | 146.36 ± 19.72 | 122.9 ± 13.48 | 140.0 ± 19.93 | **0.02** | 0.63 | - | - | - |
| | | FL | before exercise | 161.60 ± 18.91 | 149.1 ± 20.63 | 152.6 ± 18.39 | 0.34 | 0.38 | 0.49 | 0.30 | 0.70 |
| | | | after exercise | 174.80 ± 19.14 | 160.5 ± 21.84 | 163.6 ± 19.44 | 0.29 | 0.27 | 0.59 | 0.31 | 0.69 |
| | | | 60 min after exercise | 166.60 ± 19.45 | 153.8 ± 19.89 | 153.9 ± 16.99 | 0.37 | 0.26 | 0.48 | 0.21 | 0.61 |
| | | αCD | before exercise | 243.10 ± 53.80 | 208.9 ± 42.99 | 1312.7 ± 1167.16 | 0.39 | 0.39 | 0.08 | 0.06 | 0.30 |
| | | | after exercise | 252.90 ± 53.86 | 218.5 ± 43.86 | 1385.5 ± 1230.21 | 0.39 | 0.39 | 0.11 | 0.07 | 0.30 |
| | | | 60 min after exercise | 238.80 ± 50.78 | 204.7 ± 40.08 | 1271.2 ± 1124.89 | 0.37 | 0.39 | 0.09 | 0.06 | 0.30 |
| Glucose | mg/dl | Placebo | before exercise | 76.91 ± 2.39 | 81.8 ± 1.93 | 82.5 ± 2.46 | **0.03** | 0.06 | - | - | - |
| | | | after exercise | 73.18 ± 2.28 | 75.0 ± 2.17 | 74.7 ± 2.17 | 0.26 | 0.58 | - | - | - |
| | | | 60 min after exercise | 76.55 ± 1.61 | 76.6 ± 1.86 | 75.9 ± 1.85 | 0.94 | 0.67 | - | - | - |
| | | FL | before exercise | 77.70 ± 2.82 | 79.1 ± 3.69 | 79.1 ± 2.19 | 0.65 | 0.59 | 0.83 | 0.51 | 0.33 |
| | | | after exercise | 72.40 ± 2.49 | 75.3 ± 3.84 | 71.8 ± 2.54 | 0.31 | 0.72 | 0.82 | 0.95 | 0.39 |
| | | | 60 min after exercise | 77.50 ± 2.40 | 78.7 ± 1.93 | 74.7 ± 1.69 | 0.53 | 0.18 | 0.74 | 0.45 | 0.64 |

| | | | | | | | | | | | |
|---|---|---|---|---|---|---|---|---|---|---|---|
| | | | before exercise | 82.80 ± 2.32 | 82.5 ± 2.77 | 83.6 ± 2.01 | 0.87 | 0.38 | 0.09 | 0.84 | 0.73 |
| | | αCD | after exercise | 75.90 ± 2.02 | 76.7 ± 2.57 | 75.9 ± 2.51 | 0.58 | 1.00 | 0.39 | 0.62 | 0.73 |
| | | | 60 min after exercise | 76.70 ± 1.99 | 79.2 ± 2.36 | 78.3 ± 2.29 | 0.15 | 0.39 | 0.95 | 0.40 | 0.42 |
| | | | before exercise | 9.97 ± 1.70 | 10.54 ± 1.02 | 8.87 ± 1.35 | 0.77 | 0.65 | - | - | - |
| | | Placebo | after exercise | 21.54 ± 3.69 | 21.25 ± 3.23 | 19.57 ± 2.35 | 0.78 | 0.34 | - | - | - |
| | | | 60 min after exercise | 7.29 ± 0.79 | 8.14 ± 0.56 | 7.91 ± 0.79 | 0.32 | 0.60 | - | - | - |
| | | | before exercise | 8.21 ± 0.97 | 9.29 ± 1.59 | 7.62 ± 1.11 | 0.51 | 0.71 | 0.39 | 0.51 | 0.49 |
| Lactate | mg/dl | FL | after exercise | 22.78 ± 3.72 | 27.09 ± 4.41 | 19.70 ± 2.79 | **0.03** | 0.11 | 0.82 | 0.29 | 0.97 |
| | | | 60 min after exercise | 8.56 ± 0.87 | 9.02 ± 0.85 | 8.02 ± 0.87 | 0.51 | 0.57 | 0.29 | 0.39 | 0.93 |
| | | | before exercise | 8.76 ± 0.76 | 8.92 ± 1.02 | 7.52 ± 0.61 | 0.92 | 0.14 | 0.54 | 0.28 | 0.39 |
| | | αCD | after exercise | 20.28 ± 2.30 | 18.46 ± 2.31 | 16.53 ± 1.86 | 0.28 | **0.01** | 0.78 | 0.50 | 0.33 |
| | | | 60 min after exercise | 8.30 ± 0.64 | 7.70 ± 0.81 | 7.74 ± 0.57 | 0.53 | 0.41 | 0.34 | 0.66 | 0.87 |
| | | | before exercise | 178.18 ± 8.56 | 180.9 ± 7.18 | 176.5 ± 7.87 | 0.42 | 0.56 | - | - | - |
| | | Placebo | after exercise | 189.00 ± 9.80 | 193.2 ± 7.82 | 189.5 ± 8.79 | 0.29 | 0.84 | - | - | - |
| | | | 60 min after exercise | 196.18 ± 11.30 | 194.2 ± 8.92 | 185.6 ± 8.50 | 0.76 | 0.08 | - | - | - |
| | | | before exercise | 165.70 ± 7.12 | 171.9 ± 8.57 | 172.1 ± 9.01 | 0.32 | 0.47 | 0.28 | 0.43 | 0.71 |
| Lactate dehydrogenase | U/l | FL | after exercise | 176.10 ± 7.88 | 179.2 ± 7.63 | 177.6 ± 9.21 | 0.27 | 0.61 | 0.32 | 0.22 | 0.36 |
| | | | 60 min after exercise | 180.50 ± 5.99 | 185.5 ± 8.13 | 175.9 ± 7.69 | 0.44 | 0.25 | 0.25 | 0.48 | 0.41 |
| | | | before exercise | 171.70 ± 8.71 | 175.8 ± 6.11 | 196.1 ± 23.35 | 0.60 | 0.33 | 0.60 | 0.60 | 0.42 |
| | | αCD | after exercise | 184.90 ± 8.95 | 181.3 ± 5.86 | 208.3 ± 26.45 | 0.59 | 0.38 | 0.76 | 0.25 | 0.49 |
| | | | 60 min after exercise | 200.60 ± 11.93 | 212.6 ± 21.12 | 204.9 ± 24.20 | 0.61 | 0.88 | 0.79 | 0.42 | 0.44 |
| | | Placebo | before exercise | 268.18 ± 9.40 | 320.91 ± 16.88 | 276.18 ± 11.56 | **0.00** | 0.46 | - | - | - |
| | | | after exercise | 282.45 ± 13.34 | 358.91 ± 19.53 | 324.45 ± 11.89 | **0.00** | **0.00** | - | - | - |
| dROM | U.CARR | FL | before exercise | 261.60 ± 9.60 | 314.70 ± 8.53 | 285.90 ± 10.81 | **0.00** | **0.00** | 0.63 | 0.75 | 0.55 |
| | | | after exercise | 277.40 ± 10.56 | 339.10 ± 9.17 | 321.10 ± 12.97 | **0.00** | **0.00** | 0.77 | 0.39 | 0.85 |
| | | αCD | before exercise | 262.70 ± 12.85 | 313.22 ± 16.04 | 280.50 ± 13.47 | **0.00** | 0.23 | 0.73 | 0.75 | 0.81 |
| | | | after exercise | 279.30 ± 16.19 | 331.78 ± 17.91 | 316.10 ± 17.38 | **0.00** | **0.02** | 0.88 | 0.34 | 0.69 |

| | | | | | | | | | | | | | | | | | |
|---|---|---|---|---|---|---|---|---|---|---|---|---|---|---|---|---|---|
| Free fatty acid | µM | Placebo | before exercise | 80.89 | ± | 15.83 | 90.72 | ± | 22.13 | 57.54 | ± | 9.40 | 0.70 | 0.09 | - | - | - |
| | | | after exercise | 430.66 | ± | 92.97 | 498.20 | ± | 143.33 | 385.79 | ± | 119.35 | 0.66 | 0.77 | - | - | - |
| | | FL | before exercise | 70.94 | ± | 20.06 | 168.52 | ± | 56.42 | 75.67 | ± | 18.66 | **0.03** | 0.86 | 0.70 | 0.20 | 0.38 |
| | | | after exercise | 363.10 | ± | 70.25 | 480.55 | ± | 95.62 | 254.04 | ± | 48.74 | 0.15 | 0.26 | 0.57 | 0.92 | 0.34 |
| | | αCD | before exercise | 107.34 | ± | 29.36 | 123.22 | ± | 28.46 | 88.60 | ± | 15.43 | 0.52 | 0.56 | 0.43 | 0.37 | 0.10 |
| | | | after exercise | 1011.77 | ± | 299.08 | 748.87 | ± | 171.27 | 416.67 | ± | 65.60 | 0.23 | 0.08 | 0.07 | 0.27 | 0.83 |
| Growth hormone | ng/ml | Placebo | before exercise | 0.20 | ± | 0.09 | 0.47 | ± | 0.20 | 0.63 | ± | 0.34 | 0.25 | 0.26 | - | - | - |
| | | | after exercise | 8.56 | ± | 2.11 | 6.60 | ± | 1.28 | 4.71 | ± | 0.99 | 0.19 | **0.04** | - | - | - |
| | | FL | before exercise | 0.32 | ± | 0.16 | 0.24 | ± | 0.07 | 0.30 | ± | 0.13 | 0.63 | 0.90 | 0.52 | 0.32 | 0.41 |
| | | | after exercise | 5.65 | ± | 1.46 | 7.89 | ± | 2.07 | 8.16 | ± | 2.25 | 0.34 | 0.31 | 0.28 | 0.59 | 0.16 |
| | | αCD | before exercise | 1.27 | ± | 0.67 | 1.07 | ± | 0.67 | 1.24 | ± | 0.71 | 0.32 | 0.95 | 0.11 | 0.38 | 0.43 |
| | | | after exercise | 5.34 | ± | 1.54 | 5.76 | ± | 1.54 | 5.70 | ± | 1.35 | 0.25 | 0.83 | 0.24 | 0.68 | 0.56 |
| Cortisol | µg/dl | Placebo | before exercise | 9.17 | ± | 1.03 | 7.89 | ± | 0.69 | 7.29 | ± | 0.60 | 0.08 | **0.02** | - | - | - |
| | | | after exercise | 11.05 | ± | 1.51 | 10.15 | ± | 1.20 | 8.39 | ± | 0.80 | 0.38 | **0.03** | - | - | - |
| | | FL | before exercise | 11.16 | ± | 1.25 | 11.75 | ± | 1.01 | 10.04 | ± | 1.03 | 0.45 | 0.17 | 0.23 | **0.00** | **0.03** |
| | | | after exercise | 10.52 | ± | 1.03 | 11.06 | ± | 1.74 | 10.57 | ± | 0.76 | 0.73 | 0.93 | 0.78 | 0.67 | 0.06 |
| | | αCD | before exercise | 9.21 | ± | 0.96 | 11.03 | ± | 1.07 | 9.90 | ± | 0.69 | 0.24 | 0.54 | 0.98 | **0.02** | **0.01** |
| | | | after exercise | 11.24 | ± | 1.32 | 10.46 | ± | 1.35 | 10.58 | ± | 1.21 | 0.17 | 0.36 | 0.93 | 0.87 | 0.14 |
| Glucagon | pg/ml | Placebo | before exercise | 152.55 | ± | 7.12 | 132.73 | ± | 6.47 | 148.45 | ± | 6.64 | **0.00** | 0.34 | - | - | - |
| | | | after exercise | 191.91 | ± | 10.53 | 171.73 | ± | 10.13 | 179.55 | ± | 11.06 | **0.00** | **0.01** | - | - | - |
| | | FL | before exercise | 150.00 | ± | 5.33 | 137.70 | ± | 5.46 | 149.60 | ± | 2.88 | 0.08 | 0.94 | 0.78 | 0.57 | 0.88 |
| | | | after exercise | 176.70 | ± | 5.95 | 161.80 | ± | 7.80 | 165.30 | ± | 5.57 | **0.05** | 0.05 | 0.24 | 0.45 | 0.28 |
| | | αCD | before exercise | 159.60 | ± | 4.43 | 150.44 | ± | 6.34 | 163.10 | ± | 8.28 | 0.06 | 0.62 | 0.42 | 0.07 | 0.18 |
| | | | after exercise | 192.70 | ± | 4.60 | 179.67 | ± | 7.29 | 186.00 | ± | 10.76 | 0.06 | 0.51 | 0.95 | 0.56 | 0.68 |
| IL-6 | pg/ml | Placebo | before exercise | 1.02 | ± | 0.13 | 0.81 | ± | 0.09 | 0.83 | ± | 0.12 | 0.13 | 0.31 | - | - | - |
| | | | after exercise | 2.08 | ± | 0.40 | 1.64 | ± | 0.25 | 1.48 | ± | 0.24 | **0.05** | 0.10 | - | - | - |
| | | FL | before exercise | 0.62 | ± | 0.05 | 0.62 | ± | 0.04 | 0.74 | ± | 0.06 | 0.95 | 0.10 | **0.01** | 0.09 | 0.51 |

| | | | | | | | | | | | |
|---|---|---|---|---|---|---|---|---|---|---|---|
| | | | after exercise | 1.55 ± 0.42 | 1.40 ± 0.35 | 1.24 ± 0.13 | 0.71 | 0.45 | 0.37 | 0.56 | 0.41 |
| | | αCD | before exercise | 0.84 ± 0.14 | 1.00 ± 0.12 | 0.89 ± 0.07 | 0.65 | 0.73 | 0.37 | 0.22 | 0.67 |
| | | | after exercise | 1.55 ± 0.20 | 1.38 ± 0.15 | 1.38 ± 0.15 | 0.52 | 0.36 | 0.27 | 0.41 | 0.75 |
| Alanine | nmol/mL | Placebo | before exercise | 374.59 ± 32.73 | 389.48 ± 31.08 | 386.60 ± 33.46 | 0.16 | 0.37 | - | - | - |
| | | | after exercise | 437.65 ± 38.84 | 452.58 ± 37.74 | 424.30 ± 40.29 | 0.30 | 0.50 | - | - | - |
| | | FL | before exercise | 413.43 ± 19.99 | 426.59 ± 21.53 | 410.17 ± 20.24 | 0.32 | 0.84 | 0.34 | 0.35 | 0.56 |
| | | | after exercise | 468.96 ± 20.90 | 484.08 ± 25.24 | 450.17 ± 25.50 | 0.17 | 0.32 | 0.50 | 0.51 | 0.60 |
| | | αCD | before exercise | 382.92 ± 20.18 | 377.06 ± 22.27 | 380.35 ± 17.58 | 0.90 | 0.94 | 0.83 | 0.76 | 0.87 |
| | | | after exercise | 409.92 ± 26.11 | 393.54 ± 27.62 | 414.84 ± 24.14 | 0.74 | 0.89 | 0.57 | 0.24 | 0.85 |
| Alpha-amino-n-butyric acid | nmol/mL | Placebo | before exercise | 14.91 ± 0.87 | 15.69 ± 1.58 | 16.83 ± 1.97 | 0.49 | 0.17 | - | - | - |
| | | | after exercise | 14.35 ± 0.82 | 14.93 ± 1.40 | 15.32 ± 1.67 | 0.59 | 0.43 | - | - | - |
| | | FL | before exercise | 17.30 ± 2.00 | 14.97 ± 1.54 | 16.43 ± 1.82 | 0.14 | 0.42 | 0.27 | 0.75 | 0.88 |
| | | | after exercise | 16.42 ± 1.74 | 14.36 ± 1.35 | 15.64 ± 1.76 | 0.13 | 0.46 | 0.28 | 0.78 | 0.90 |
| | | αCD | before exercise | 17.67 ± 1.51 | 19.69 ± 1.57 | 17.92 ± 1.67 | **0.04** | 0.87 | 0.12 | 0.09 | 0.68 |
| | | | after exercise | 16.06 ± 1.39 | 18.04 ± 1.41 | 16.71 ± 1.48 | **0.04** | 0.67 | 0.29 | 0.14 | 0.54 |
| Arginine | nmol/mL | Placebo | before exercise | 87.82 ± 6.00 | 97.13 ± 5.29 | 95.62 ± 6.54 | **0.04** | 0.20 | - | - | - |
| | | | after exercise | 94.08 ± 5.19 | 103.41 ± 6.44 | 97.15 ± 6.09 | 0.13 | 0.61 | - | - | - |
| | | FL | before exercise | 97.27 ± 5.93 | 103.10 ± 6.52 | 97.55 ± 7.26 | 0.28 | 0.97 | 0.28 | 0.48 | 0.84 |
| | | | after exercise | 105.53 ± 7.85 | 100.27 ± 6.22 | 105.46 ± 7.32 | 0.24 | 0.99 | 0.23 | 0.73 | 0.39 |
| | | αCD | before exercise | 91.63 ± 7.33 | 90.11 ± 3.57 | 91.23 ± 4.65 | 0.94 | 0.96 | 0.69 | 0.31 | 0.60 |
| | | | after exercise | 94.63 ± 4.19 | 96.29 ± 3.88 | 93.41 ± 3.78 | 0.52 | 0.82 | 0.94 | 0.38 | 0.62 |
| Asparagine | nmol/mL | Placebo | before exercise | 53.12 ± 2.23 | 56.40 ± 2.72 | 58.13 ± 2.95 | **0.05** | **0.01** | - | - | - |
| | | | after exercise | 53.29 ± 2.55 | 56.11 ± 2.37 | 53.78 ± 2.77 | 0.17 | 0.82 | - | - | - |
| | | FL | before exercise | 58.43 ± 2.15 | 56.86 ± 1.85 | 58.95 ± 2.67 | 0.49 | 0.82 | 0.10 | 0.89 | 0.84 |
| | | | after exercise | 58.37 ± 3.00 | 55.50 ± 2.57 | 57.33 ± 3.84 | 0.12 | 0.66 | 0.21 | 0.86 | 0.46 |
| | | αCD | before exercise | 56.09 ± 1.50 | 56.19 ± 2.97 | 57.40 ± 2.41 | 0.96 | 0.66 | 0.29 | 0.96 | 0.85 |
| | | | after exercise | 53.94 ± 1.47 | 54.70 ± 2.64 | 56.00 ± 1.88 | 0.82 | 0.33 | 0.83 | 0.70 | 0.52 |

| | | | | | | | | | | | |
|---|---|---|---|---|---|---|---|---|---|---|---|
| Aspartic acid | nmol/mL | Placebo | before exercise | 3.45 ± 0.37 | 4.75 ± 0.56 | 3.43 ± 0.25 | **0.04** | 0.97 | - | - | - |
| | | | after exercise | 5.78 ± 1.20 | 6.64 ± 1.09 | 4.49 ± 0.47 | 0.71 | 0.25 | - | - | - |
| | | FL | before exercise | 6.09 ± 1.31 | 6.43 ± 1.35 | 7.97 ± 1.30 | 0.91 | 0.09 | **0.05** | 0.23 | **0.00** |
| | | | after exercise | 7.42 ± 1.77 | 6.50 ± 1.18 | 7.41 ± 2.18 | 0.48 | 1.00 | 0.45 | 0.93 | 0.19 |
| | | αCD | before exercise | 5.66 ± 1.11 | 4.60 ± 0.43 | 5.98 ± 0.81 | 0.21 | 0.98 | 0.06 | 0.84 | **0.00** |
| | | | after exercise | 6.80 ± 1.42 | 6.73 ± 0.93 | 5.54 ± 0.70 | 0.63 | 0.28 | 0.59 | 0.95 | 0.22 |
| Beta-alanine | nmol/mL | Placebo | before exercise | 4.57 ± 0.34 | 5.93 ± 0.46 | 7.66 ± 0.41 | **0.03** | **0.00** | - | - | - |
| | | | after exercise | 4.82 ± 0.34 | 6.08 ± 0.47 | 7.55 ± 0.46 | **0.05** | **0.00** | - | - | - |
| | | FL | before exercise | 6.49 ± 0.34 | 6.94 ± 0.47 | 9.43 ± 0.44 | 0.32 | **0.00** | **0.00** | 0.14 | **0.01** |
| | | | after exercise | 6.68 ± 0.53 | 6.62 ± 0.59 | 9.13 ± 0.45 | 0.91 | **0.00** | **0.01** | 0.48 | **0.02** |
| | | αCD | before exercise | 5.88 ± 0.61 | 6.36 ± 0.37 | 7.88 ± 0.58 | 0.51 | **0.02** | 0.07 | 0.49 | 0.76 |
| | | | after exercise | 6.12 ± 0.49 | 6.11 ± 0.40 | 7.70 ± 0.60 | 0.85 | 0.07 | **0.04** | 0.96 | 0.85 |
| Beta-amino isobutyric acid | nmol/mL | Placebo | before exercise | 2.52 ± 0.55 | 2.73 ± 0.71 | 3.17 ± 0.27 | 0.27 | 0.62 | - | - | - |
| | | | after exercise | 2.60 ± 0.52 | 2.90 ± 0.79 | 3.23 ± 0.48 | 0.10 | 0.26 | - | - | - |
| | | FL | before exercise | 3.02 ± 0.50 | 3.24 ± 0.38 | 3.28 ± 0.43 | 0.28 | 0.37 | 0.52 | 0.51 | 0.86 |
| | | | after exercise | 2.80 ± 0.54 | 3.30 ± 0.39 | 3.35 ± 0.53 | 0.10 | 0.27 | 0.80 | 0.63 | 0.89 |
| | | αCD | before exercise | 3.05 ± 0.23 | 2.95 ± 0.41 | 3.45 ± 0.28 | 0.46 | 0.07 | 0.36 | 0.78 | 0.51 |
| | | | after exercise | 3.05 ± 0.27 | 2.97 ± 0.42 | 3.38 ± 0.23 | 0.14 | 0.77 | 0.46 | 0.94 | 0.78 |
| Citrulline | nmol/mL | Placebo | before exercise | 32.15 ± 1.85 | 35.40 ± 2.28 | 36.22 ± 2.11 | **0.02** | **0.02** | - | - | - |
| | | | after exercise | 31.89 ± 1.43 | 34.07 ± 1.58 | 33.33 ± 1.46 | **0.02** | 0.13 | - | - | - |
| | | FL | before exercise | 32.91 ± 1.52 | 31.30 ± 1.50 | 31.69 ± 1.63 | 0.25 | 0.47 | 0.76 | 0.16 | 0.11 |
| | | | after exercise | 31.03 ± 1.44 | 29.66 ± 1.05 | 30.30 ± 1.54 | 0.28 | 0.49 | 0.68 | **0.03** | 0.17 |
| | | αCD | before exercise | 31.73 ± 1.41 | 33.12 ± 1.21 | 31.05 ± 1.32 | 0.24 | 0.67 | 0.86 | 0.42 | 0.06 |
| | | | after exercise | 30.23 ± 1.64 | 32.62 ± 1.83 | 31.01 ± 1.24 | 0.14 | 0.46 | 0.45 | 0.55 | 0.25 |
| Cystine | nmol/mL | Placebo | before exercise | 36.86 ± 1.67 | 40.82 ± 1.32 | 39.93 ± 1.79 | **0.03** | 0.23 | - | - | - |
| | | | after exercise | 37.13 ± 1.98 | 42.88 ± 1.67 | 39.85 ± 1.66 | **0.01** | 0.32 | - | - | - |
| | | FL | before exercise | 37.85 ± 1.76 | 41.49 ± 2.14 | 41.53 ± 2.15 | **0.04** | **0.04** | 0.69 | 0.79 | 0.57 |

| | | | | | | | | | | | |
|---|---|---|---|---|---|---|---|---|---|---|---|
| | | | after exercise | 36.53 ± 1.91 | 43.24 ± 2.02 | 43.06 ± 2.18 | **0.00** | **0.01** | 0.83 | 0.89 | 0.25 |
| | | αCD | before exercise | 37.14 ± 2.96 | 42.46 ± 2.93 | 42.34 ± 2.48 | 0.06 | **0.05** | 0.93 | 0.59 | 0.43 |
| | | | after exercise | 37.58 ± 3.19 | 44.90 ± 2.41 | 41.75 ± 2.04 | **0.01** | 0.09 | 0.90 | 0.49 | 0.47 |
| Glutamic acid | nmol/mL | Placebo | before exercise | 41.05 ± 3.55 | 53.00 ± 5.15 | 43.16 ± 3.79 | **0.04** | 0.70 | - | - | - |
| | | | after exercise | 67.31 ± 9.51 | 65.26 ± 7.87 | 58.64 ± 5.87 | 0.85 | 0.33 | - | - | - |
| | | FL | before exercise | 54.88 ± 7.85 | 65.48 ± 12.83 | 64.38 ± 15.83 | 0.24 | 0.46 | 0.11 | 0.36 | 0.19 |
| | | | after exercise | 84.62 ± 13.88 | 77.52 ± 13.14 | 76.35 ± 14.06 | 0.42 | 0.52 | 0.31 | 0.42 | 0.24 |
| | | αCD | before exercise | 62.16 ± 8.52 | 53.01 ± 6.04 | 63.20 ± 9.88 | 0.17 | 0.93 | **0.03** | 1.00 | 0.06 |
| | | | after exercise | 87.16 ± 14.50 | 73.92 ± 9.08 | 67.45 ± 9.01 | 0.15 | 0.08 | 0.26 | 0.48 | 0.41 |
| Glutamine | nmol/mL | Placebo | before exercise | 562.26 ± 20.89 | 572.32 ± 20.86 | 594.17 ± 20.59 | 0.40 | **0.04** | - | - | - |
| | | | after exercise | 566.38 ± 21.29 | 590.99 ± 22.68 | 568.12 ± 23.80 | 0.14 | 0.91 | - | - | - |
| | | FL | before exercise | 571.25 ± 15.62 | 567.44 ± 21.28 | 583.52 ± 23.11 | 0.84 | 0.46 | 0.74 | 0.87 | 0.73 |
| | | | after exercise | 574.73 ± 20.19 | 581.08 ± 12.57 | 590.40 ± 22.16 | 0.70 | 0.42 | 0.78 | 0.71 | 0.50 |
| | | αCD | before exercise | 548.43 ± 21.13 | 548.94 ± 16.73 | 557.27 ± 20.54 | 0.50 | 0.68 | 0.65 | 0.41 | 0.22 |
| | | | after exercise | 532.25 ± 22.20 | 545.33 ± 14.80 | 566.55 ± 14.98 | 0.32 | 0.12 | 0.28 | 0.13 | 0.96 |
| Glycine | nmol/mL | Placebo | before exercise | 275.54 ± 20.76 | 287.04 ± 24.56 | 286.75 ± 24.94 | 0.23 | 0.37 | - | - | - |
| | | | after exercise | 274.11 ± 22.17 | 287.17 ± 25.10 | 269.81 ± 26.80 | 0.18 | 0.73 | - | - | - |
| | | FL | before exercise | 268.85 ± 7.90 | 258.07 ± 6.76 | 259.62 ± 9.24 | 0.09 | 0.21 | 0.78 | 0.29 | 0.34 |
| | | | after exercise | 268.33 ± 12.34 | 254.11 ± 9.29 | 253.44 ± 13.41 | 0.12 | 0.08 | 0.83 | 0.25 | 0.60 |
| | | αCD | before exercise | 253.79 ± 7.89 | 249.42 ± 17.74 | 245.70 ± 8.94 | 0.85 | 0.47 | 0.36 | 0.25 | 0.15 |
| | | | after exercise | 241.57 ± 8.00 | 242.03 ± 16.54 | 240.21 ± 7.99 | 0.96 | 0.89 | 0.20 | 0.17 | 0.32 |
| Histidine | nmol/mL | Placebo | before exercise | 81.81 ± 2.81 | 83.05 ± 1.86 | 89.57 ± 3.04 | 0.59 | **0.01** | - | - | - |
| | | | after exercise | 83.17 ± 3.05 | 86.13 ± 2.82 | 86.26 ± 3.57 | 0.34 | 0.34 | - | - | - |
| | | FL | before exercise | 84.88 ± 1.68 | 84.08 ± 2.60 | 87.10 ± 1.66 | 0.79 | 0.21 | 0.37 | 0.75 | 0.50 |
| | | | after exercise | 86.36 ± 2.71 | 83.60 ± 3.34 | 86.51 ± 2.13 | 0.31 | 0.94 | 0.45 | 0.57 | 0.95 |
| | | αCD | before exercise | 85.62 ± 1.62 | 84.99 ± 2.45 | 89.15 ± 2.26 | 0.81 | 0.23 | 0.27 | 0.53 | 0.91 |
| | | | after exercise | 84.43 ± 1.94 | 84.88 ± 2.15 | 88.59 ± 1.87 | 1.00 | 0.18 | 0.74 | 0.74 | 0.58 |

| | | | | | | | | | | |
|---|---|---|---|---|---|---|---|---|---|---|
| Hy proline | nmol/mL | Placebo | before exercise | 10.86 ± 1.16 | 10.56 ± 1.36 | 10.04 ± 1.33 | 0.73 | 0.37 | - | - | - |
| | | | after exercise | 10.24 ± 0.92 | 9.95 ± 1.17 | 8.88 ± 1.06 | 0.72 | 0.11 | - | - | - |
| | | FL | before exercise | 13.52 ± 1.89 | 9.90 ± 1.08 | 11.96 ± 2.43 | 0.06 | 0.38 | 0.24 | 0.71 | 0.49 |
| | | | after exercise | 13.03 ± 1.70 | 9.32 ± 0.99 | 10.89 ± 2.05 | **0.04** | 0.17 | 0.15 | 0.69 | 0.38 |
| | | αCD | before exercise | 12.33 ± 1.09 | 14.73 ± 2.21 | 12.07 ± 1.53 | 0.27 | 0.81 | 0.37 | 0.11 | 0.33 |
| | | | after exercise | 11.09 ± 1.01 | 13.16 ± 1.99 | 11.33 ± 1.49 | 0.27 | 0.83 | 0.54 | 0.16 | 0.19 |
| Isoleucine | nmol/mL | Placebo | before exercise | 52.95 ± 2.70 | 57.45 ± 3.07 | 61.72 ± 4.73 | 0.09 | **0.05** | - | - | - |
| | | | after exercise | 57.17 ± 2.54 | 60.17 ± 3.58 | 59.99 ± 4.00 | 0.35 | 0.27 | - | - | - |
| | | FL | before exercise | 60.88 ± 2.64 | 60.79 ± 3.02 | 66.75 ± 4.28 | 0.98 | 0.13 | **0.05** | 0.45 | 0.44 |
| | | | after exercise | 61.43 ± 3.11 | 62.80 ± 2.82 | 67.13 ± 3.89 | 0.58 | 0.08 | 0.30 | 0.58 | 0.22 |
| | | αCD | before exercise | 59.58 ± 2.27 | 68.10 ± 3.95 | 69.29 ± 4.65 | 0.06 | 0.11 | 0.08 | **0.04** | 0.27 |
| | | | after exercise | 60.96 ± 2.62 | 68.06 ± 3.81 | 68.13 ± 2.76 | 0.06 | **0.02** | 0.31 | 0.15 | 0.12 |
| Leucine | nmol/mL | Placebo | before exercise | 99.25 ± 3.60 | 108.20 ± 4.23 | 117.30 ± 7.16 | **0.05** | **0.02** | - | - | - |
| | | | after exercise | 107.50 ± 3.54 | 114.15 ± 5.74 | 114.53 ± 6.29 | 0.25 | 0.15 | - | - | - |
| | | FL | before exercise | 120.13 ± 4.42 | 117.62 ± 6.36 | 123.52 ± 5.49 | 0.75 | 0.54 | **0.00** | 0.23 | 0.51 |
| | | | after exercise | 122.48 ± 6.06 | 121.05 ± 6.59 | 125.87 ± 6.63 | 0.79 | 0.53 | **0.04** | 0.44 | 0.23 |
| | | αCD | before exercise | 117.92 ± 4.20 | 132.40 ± 6.32 | 132.99 ± 5.75 | **0.02** | 0.07 | **0.00** | **0.00** | 0.11 |
| | | | after exercise | 121.57 ± 5.89 | 133.78 ± 7.20 | 132.09 ± 4.47 | **0.01** | **0.03** | 0.05 | **0.04** | **0.04** |
| Lysine | nmol/mL | Placebo | before exercise | 158.31 ± 7.68 | 167.10 ± 8.54 | 184.01 ± 11.64 | 0.19 | **0.02** | - | - | - |
| | | | after exercise | 169.80 ± 8.05 | 179.38 ± 8.05 | 183.34 ± 12.60 | 0.28 | 0.23 | - | - | - |
| | | FL | before exercise | 188.99 ± 4.94 | 181.94 ± 5.77 | 189.39 ± 3.51 | 0.34 | 0.93 | **0.00** | 0.17 | 0.68 |
| | | | after exercise | 198.97 ± 6.66 | 185.07 ± 5.02 | 194.78 ± 6.96 | **0.03** | 0.22 | **0.01** | 0.57 | 0.45 |
| | | αCD | before exercise | 180.76 ± 7.02 | 186.59 ± 3.07 | 187.29 ± 8.50 | 0.55 | 0.55 | **0.05** | 0.06 | 0.83 |
| | | | after exercise | 182.11 ± 7.22 | 187.52 ± 5.94 | 186.51 ± 5.89 | 0.60 | 0.63 | 0.27 | 0.44 | 0.83 |
| Methionine | nmol/mL | Placebo | before exercise | 23.15 ± 0.92 | 25.10 ± 1.08 | 26.75 ± 1.67 | **0.03** | **0.04** | - | - | - |
| | | | after exercise | 26.19 ± 1.41 | 27.65 ± 1.11 | 27.42 ± 1.46 | 0.37 | 0.48 | - | - | - |
| | | FL | before exercise | 27.00 ± 1.20 | 26.74 ± 1.28 | 26.19 ± 1.50 | 0.86 | 0.56 | **0.02** | 0.34 | 0.81 |

| | | | | | | | | | | | |
|---|---|---|---|---|---|---|---|---|---|---|---|
| | | | after exercise | 29.45 ± 1.98 | 28.00 ± 1.33 | 28.37 ± 2.44 | 0.21 | 0.45 | 0.19 | 0.84 | 0.74 |
| | | αCD | before exercise | 26.14 ± 1.36 | 27.81 ± 1.21 | 27.31 ± 1.30 | 0.34 | 0.54 | 0.08 | 0.11 | 0.80 |
| | | | after exercise | 28.11 ± 1.29 | 29.33 ± 1.53 | 27.92 ± 1.27 | 0.50 | 0.85 | 0.33 | 0.37 | 0.80 |
| Mono ethanolamine | nmol/mL | Placebo | before exercise | 8.79 ± 0.33 | 9.31 ± 0.46 | 8.73 ± 0.45 | 0.18 | 0.87 | - | - | - |
| | | | after exercise | 10.53 ± 0.48 | 10.49 ± 0.73 | 9.68 ± 0.43 | 0.96 | 0.07 | - | - | - |
| | | FL | before exercise | 10.66 ± 0.64 | 10.62 ± 0.62 | 9.32 ± 0.54 | 0.94 | **0.03** | **0.02** | 0.10 | 0.41 |
| | | | after exercise | 11.93 ± 1.01 | 11.36 ± 0.93 | 10.37 ± 0.69 | 0.42 | **0.04** | 0.21 | 0.47 | 0.40 |
| | | αCD | before exercise | 10.36 ± 0.42 | 9.63 ± 0.39 | 9.67 ± 0.48 | **0.01** | **0.04** | **0.01** | 0.61 | 0.17 |
| | | | after exercise | 11.50 ± 0.78 | 10.72 ± 0.60 | 10.62 ± 0.61 | **0.01** | 0.11 | 0.29 | 0.82 | 0.22 |
| Ornithine | nmol/mL | Placebo | before exercise | 58.02 ± 3.55 | 59.58 ± 3.30 | 63.92 ± 3.36 | 0.30 | 0.10 | - | - | - |
| | | | after exercise | 52.28 ± 2.83 | 57.16 ± 2.98 | 55.64 ± 3.57 | 0.14 | 0.35 | - | - | - |
| | | FL | before exercise | 60.32 ± 4.34 | 54.77 ± 3.87 | 62.87 ± 3.66 | 0.05 | 0.55 | 0.68 | 0.35 | 0.83 |
| | | | after exercise | 56.83 ± 4.14 | 51.28 ± 2.11 | 53.11 ± 4.24 | 0.08 | 0.24 | 0.37 | 0.13 | 0.65 |
| | | αCD | before exercise | 55.37 ± 3.13 | 57.48 ± 2.47 | 60.28 ± 2.87 | 0.32 | 0.28 | 0.59 | 0.63 | 0.43 |
| | | | after exercise | 47.86 ± 2.83 | 51.10 ± 2.02 | 53.02 ± 2.35 | 0.39 | 0.21 | 0.28 | 0.13 | 0.56 |
| Phenylalanine | nmol/mL | Placebo | before exercise | 52.85 ± 1.60 | 57.47 ± 2.70 | 58.27 ± 2.11 | **0.03** | **0.01** | - | - | - |
| | | | after exercise | 54.83 ± 2.02 | 58.83 ± 3.28 | 57.25 ± 2.16 | 0.16 | 0.22 | - | - | - |
| | | FL | before exercise | 61.47 ± 2.46 | 61.80 ± 1.92 | 59.50 ± 1.80 | 0.91 | 0.23 | **0.01** | 0.21 | 0.67 |
| | | | after exercise | 61.74 ± 2.45 | 60.84 ± 2.23 | 60.60 ± 3.14 | 0.74 | 0.61 | **0.04** | 0.62 | 0.38 |
| | | αCD | before exercise | 61.24 ± 2.67 | 64.61 ± 3.30 | 62.09 ± 2.93 | 0.25 | 0.71 | **0.01** | 0.11 | 0.30 |
| | | | after exercise | 61.17 ± 2.37 | 65.38 ± 3.10 | 61.48 ± 2.48 | 0.11 | 0.83 | 0.05 | 0.17 | 0.21 |
| Proline | nmol/mL | Placebo | before exercise | 189.63 ± 28.74 | 190.04 ± 23.87 | 207.05 ± 41.76 | 0.95 | 0.29 | - | - | - |
| | | | after exercise | 198.15 ± 26.07 | 203.16 ± 29.39 | 203.47 ± 37.96 | 0.41 | 0.73 | - | - | - |
| | | FL | before exercise | 171.06 ± 12.26 | 167.77 ± 11.03 | 179.90 ± 16.07 | 0.70 | 0.43 | 0.57 | 0.42 | 0.57 |
| | | | after exercise | 178.57 ± 12.20 | 171.66 ± 10.36 | 183.14 ± 17.25 | 0.09 | 0.58 | 0.52 | 0.34 | 0.64 |
| | | αCD | before exercise | 162.03 ± 8.96 | 163.19 ± 11.69 | 166.26 ± 11.12 | 0.85 | 0.57 | 0.39 | 0.36 | 0.38 |
| | | | after exercise | 162.71 ± 8.29 | 163.49 ± 11.81 | 166.10 ± 7.85 | 0.91 | 0.65 | 0.23 | 0.26 | 0.37 |

| | | | | | | | | | | | |
|---|---|---|---|---|---|---|---|---|---|---|---|
| Serine | nmol/mL | Placebo | before exercise | 122.65 ± 5.53 | 129.82 ± 5.87 | 131.78 ± 4.77 | **0.04** | **0.01** | - | - | - |
| | | | after exercise | 130.35 ± 6.29 | 135.78 ± 4.89 | 129.97 ± 4.57 | 0.17 | 0.94 | - | - | - |
| | | FL | before exercise | 128.74 ± 5.63 | 126.30 ± 4.98 | 126.28 ± 5.31 | 0.62 | 0.63 | 0.45 | 0.66 | 0.45 |
| | | | after exercise | 137.58 ± 7.55 | 129.02 ± 5.13 | 131.71 ± 8.39 | 0.11 | 0.44 | 0.47 | 0.35 | 0.85 |
| | | αCD | before exercise | 114.67 ± 4.70 | 120.36 ± 4.66 | 116.36 ± 4.54 | 0.56 | 0.80 | 0.29 | 0.24 | **0.03** |
| | | | after exercise | 117.18 ± 4.78 | 124.24 ± 6.55 | 116.69 ± 4.59 | 0.35 | 0.91 | 0.12 | 0.17 | 0.05 |
| Taurine | nmol/ml | Placebo | before exercise | 57.62 ± 7.65 | 74.69 ± 8.30 | 56.97 ± 5.58 | **0.04** | 0.87 | - | - | - |
| | | | after exercise | 89.96 ± 18.01 | 91.12 ± 11.82 | 66.80 ± 7.11 | 0.94 | 0.14 | - | - | - |
| | | FL | before exercise | 80.34 ± 12.33 | 91.77 ± 11.90 | 87.38 ± 11.58 | 0.41 | 0.61 | 0.13 | 0.25 | **0.02** |
| | | | after exercise | 104.49 ± 16.68 | 97.57 ± 12.19 | 106.03 ± 17.51 | 0.60 | 0.93 | 0.56 | 0.71 | **0.04** |
| | | αCD | before exercise | 82.36 ± 11.86 | 62.34 ± 4.17 | 75.37 ± 6.55 | 0.06 | 0.55 | 0.09 | 0.23 | **0.04** |
| | | | after exercise | 97.61 ± 17.55 | 91.46 ± 10.73 | 85.42 ± 12.34 | 0.49 | 0.42 | 0.77 | 0.98 | 0.20 |
| Threonine | nmol/mL | Placebo | before exercise | 128.26 ± 8.52 | 141.13 ± 13.68 | 144.33 ± 11.56 | 0.10 | **0.02** | - | - | - |
| | | | after exercise | 130.94 ± 7.96 | 141.23 ± 12.35 | 137.64 ± 11.26 | 0.19 | 0.37 | - | - | - |
| | | FL | before exercise | 143.96 ± 7.26 | 137.80 ± 6.14 | 137.16 ± 6.36 | 0.41 | 0.37 | 0.18 | 0.83 | 0.60 |
| | | | after exercise | 143.84 ± 5.56 | 136.86 ± 7.68 | 134.93 ± 7.39 | 0.30 | 0.16 | 0.21 | 0.77 | 0.85 |
| | | αCD | before exercise | 128.66 ± 4.43 | 132.70 ± 5.48 | 130.07 ± 4.42 | 0.37 | 0.80 | 0.97 | 0.60 | 0.28 |
| | | | after exercise | 124.84 ± 3.91 | 129.63 ± 6.07 | 128.13 ± 3.52 | 0.37 | 0.57 | 0.51 | 0.44 | 0.45 |
| Tryptophan | nmol/mL | Placebo | before exercise | 49.15 ± 1.45 | 51.48 ± 1.77 | 52.96 ± 2.14 | 0.25 | **0.03** | - | - | - |
| | | | after exercise | 47.25 ± 1.98 | 49.93 ± 1.89 | 48.66 ± 2.41 | 0.19 | 0.58 | - | - | - |
| | | FL | before exercise | 54.46 ± 2.31 | 50.80 ± 1.82 | 47.91 ± 1.68 | 0.07 | **0.01** | 0.06 | 0.79 | 0.08 |
| | | | after exercise | 49.90 ± 1.91 | 47.87 ± 1.71 | 46.74 ± 2.11 | 0.23 | 0.14 | 0.35 | 0.43 | 0.56 |
| | | αCD | before exercise | 48.80 ± 1.69 | 53.76 ± 2.89 | 52.97 ± 1.82 | 0.08 | 0.07 | 0.87 | 0.49 | 1.00 |
| | | | after exercise | 41.80 ± 2.40 | 47.81 ± 3.15 | 47.45 ± 1.68 | **0.05** | 0.07 | 0.09 | 0.56 | 0.69 |
| Tyrosine | nmol/mL | Placebo | before exercise | 51.68 ± 2.19 | 54.08 ± 2.02 | 56.95 ± 2.75 | 0.21 | 0.08 | - | - | - |
| | | | after exercise | 56.18 ± 2.16 | 57.85 ± 2.53 | 56.03 ± 2.48 | 0.43 | 0.94 | - | - | - |
| | | FL | before exercise | 62.80 ± 3.11 | 60.62 ± 2.42 | 59.03 ± 2.92 | 0.51 | 0.13 | **0.01** | 0.05 | 0.61 |

| | | | | | | | | | | | |
|---|---|---|---|---|---|---|---|---|---|---|---|
| | | | after exercise | 65.32 ± 2.87 | 62.69 ± 2.69 | 61.69 ± 3.49 | 0.25 | 0.10 | **0.02** | 0.20 | 0.20 |
| | | αCD | before exercise | 60.14 ± 2.48 | 67.29 ± 4.46 | 65.09 ± 3.68 | 0.06 | 0.24 | **0.02** | **0.01** | 0.09 |
| | | | after exercise | 62.44 ± 2.53 | 68.16 ± 4.26 | 64.03 ± 2.57 | 0.10 | 0.44 | 0.07 | **0.04** | **0.04** |
| Valine | nmol/mL | Placebo | before exercise | 193.76 ± 6.25 | 208.12 ± 7.16 | 220.42 ± 11.32 | **0.03** | **0.01** | - | - | - |
| | | | after exercise | 197.07 ± 6.42 | 210.24 ± 9.55 | 211.91 ± 11.17 | 0.13 | 0.10 | - | - | - |
| | | FL | before exercise | 230.42 ± 7.55 | 224.58 ± 5.64 | 228.64 ± 6.68 | 0.61 | 0.78 | **0.00** | 0.09 | 0.55 |
| | | | after exercise | 225.76 ± 6.65 | 222.36 ± 6.86 | 225.23 ± 6.75 | 0.69 | 0.91 | **0.01** | 0.32 | 0.33 |
| | | αCD | before exercise | 224.46 ± 7.08 | 255.82 ± 13.53 | 245.48 ± 10.97 | **0.01** | 0.08 | **0.00** | **0.00** | 0.13 |
| | | | after exercise | 220.79 ± 8.84 | 247.66 ± 13.87 | 237.87 ± 8.68 | **0.01** | 0.06 | **0.04** | **0.03** | 0.09 |
| 1-Methyl L-histidine | nmol/mL | Placebo | before exercise | 4.23 ± 0.87 | 4.66 ± 0.76 | 6.45 ± 0.75 | 0.45 | 0.09 | - | - | - |
| | | | after exercise | 4.33 ± 0.85 | 4.78 ± 0.78 | 6.28 ± 0.75 | 0.58 | 0.15 | - | - | - |
| | | FL | before exercise | 6.17 ± 1.56 | 4.82 ± 1.08 | 6.27 ± 1.33 | 0.47 | 0.58 | 0.31 | 0.90 | 0.91 |
| | | | after exercise | 5.92 ± 1.48 | 4.93 ± 0.96 | 5.90 ± 1.16 | 0.48 | 0.71 | 0.40 | 0.90 | 0.78 |
| | | αCD | before exercise | 5.47 ± 1.29 | 7.99 ± 3.33 | 6.65 ± 1.30 | 0.50 | 0.54 | 0.46 | 0.30 | 0.89 |
| | | | after exercise | 5.44 ± 1.33 | 7.26 ± 2.89 | 6.29 ± 1.10 | 0.55 | 0.87 | 0.52 | 0.40 | 1.00 |
| 3-Methyl histidine | nmol/mL | Placebo | before exercise | 4.53 ± 0.30 | 4.70 ± 0.29 | 4.75 ± 0.19 | 0.64 | 0.38 | - | - | - |
| | | | after exercise | 4.45 ± 0.27 | 4.55 ± 0.33 | 4.65 ± 0.23 | 0.74 | 0.49 | - | - | - |
| | | FL | before exercise | 5.36 ± 0.37 | 5.41 ± 0.40 | 5.23 ± 0.25 | 0.88 | 0.60 | 0.09 | 0.16 | 0.15 |
| | | | after exercise | 5.30 ± 0.42 | 5.25 ± 0.36 | 5.30 ± 0.27 | 0.88 | 1.00 | 0.10 | 0.17 | 0.08 |
| | | αCD | before exercise | 4.88 ± 0.16 | 5.87 ± 0.56 | 5.48 ± 0.29 | 0.18 | 0.06 | 0.32 | 0.07 | **0.05** |
| | | | after exercise | 4.94 ± 0.30 | 5.74 ± 0.57 | 5.41 ± 0.27 | 0.20 | **0.05** | 0.24 | 0.07 | **0.05** |

All values are expressed as mean ± standard error.

1) two-tailed paired *t*-test

2) two-tailed unpaired *t*-test

Supplementary Table 6. RPE at baseline, 4 and 8 weeks in human study.

| | Group | Time point (min) | Baseline | 4 weeks | 8 weeks | *P*-value (vs. baseline)[1] | | *P*-value (vs. placebo)[2] | | |
|---|---|---|---|---|---|---|---|---|---|---|
| | | | | | | 4 weeks | 8 weeks | Baseline | 4 weeks | 8 weeks |
| RPE | Placebo | 0 | 8.9 ± 0.6 | 9.0 ± 0.6 | 9.2 ± 0.6 | 0.75 | 0.52 | - | - | - |
| | | 50 | 16.2 ± 0.6 | 15.6 ± 0.6 | 15.3 ± 0.6 | 0.11 | **0.01** | - | - | - |
| | FL | 0 | 6.8 ± 0.3 | 7.0 ± 0.3 | 7.2 ± 0.6 | 0.59 | 0.85 | **0.01** | **0.02** | **0.02** |
| | | 50 | 15.4 ± 0.9 | 15.0 ± 0.8 | 14.8 ± 0.8 | 0.28 | 0.08 | 0.39 | 0.48 | 0.41 |
| | αCD | 0 | 8.3 ± 0.6 | 8.0 ± 0.5 | 7.4 ± 0.5 | 0.46 | 0.13 | 0.47 | 0.27 | **0.03** |
| | | 50 | 16.6 ± 0.7 | 16.1 ± 0.3 | 14.3 ± 0.6 | 0.32 | **0.01** | 0.78 | 0.50 | 0.27 |

Mean ± standard error

1) Wilcoxon signed-rank test

2) Mann-Whitney *U*-test

Supplementary Table 7. Muscle mass at baseline, 4 and 8 weeks in human study.

| | Group | Baseline | 4 weeks | 8 weeks | *P*-value (vs. baseline)[1) ] | | *P*-value (vs. placebo)[2)] | | |
|---|---|---|---|---|---|---|---|---|---|
| | | | | | 4 weeks | 8 weeks | Baseline | 4 weeks | 8 weeks |
| Muscle mass (kg) | Placebo | 48.66 ± 1.97 | 48.85 ± 1.99 | 48.74 ± 1.94 | 0.27 | 0.75 | - | - | - |
| | FL | 50.39 ± 1.27 | 50.02 ± 1.21 | 50.04 ± 1.09 | 0.19 | 0.40 | 0.48 | 0.63 | 0.58 |
| | αCD | 52.42 ± 1.64 | 52.49 ± 1.74 | 52.48 ± 1.64 | 0.74 | 0.83 | 0.16 | 0.19 | 0.16 |
| Body fat mass (kg) | Placebo | 12.39 ± 1.52 | 12.54 ± 1.64 | 12.35 ± 1.69 | 0.48 | 0.85 | - | - | - |
| | FL | 9.75 ± 1.41 | 9.71 ± 1.24 | 9.83 ± 1.22 | 0.87 | 0.79 | 0.22 | 0.19 | 0.25 |
| | αCD | 12.55 ± 2.02 | 12.90 ± 2.18 | 13.70 ± 2.30 | 0.17 | **0.02** | 0.95 | 0.89 | 0.64 |

mean ± standard error

1) two-tailed paired *t*-test

2) two-tailed unpaired *t*-test

Supplementary Table 8. Primers and probes used for quantitative real-time PCR for *Bacteroides* species.

| Target | Primer name | Sequence(5'-3') | Probe name | Sequence(5'Fam-3'Tam) | Reference |
|---|---|---|---|---|---|
| *B. caccae* | B.cac-TaqMan-F | AAACCCATACGCCGCAAG | B.cac-TaqMan-Prb | TGTGAAGGTGCTGCATGGTTGTCGT | 39) |
| | B.cac-TaqMan-R | GACACCTCACGGCACGAG | | | |
| *B. coprocola* | BaCOP-F | TATGGTGAGATTGCATGATGG | - | | 40) |
| | BaCOP-R | ATGAACGTCAGTTACAGTTTAGCAA | - | | |
| *B. coprophilus* | BaCPP-F | GGGTTGTAAACTTCTTTTGTGC | - | | 40) |
| | BaCPP-R | GCCTCAACCGTACTCAAGGT | - | | |
| *B. dorei* | BaDOR-F | GGAAACGGTTCAGCTAGCAATA | - | | 40) |
| | BaDOR-R | AGTCTTGTCAGAGTCCTCAGCATC | - | | |
| *B. eggerthii* | B.egg-TaqMan-F | CCCGATAGTATAGTTTTTCCGC | B.egg-TaqMan-Prb | TTCGGTTATCGATGGGGATGCGTTC | 39) |
| | B.egg-TaqMan-R | TCCTCTCAGAACCCCTATCCAT | | | |
| *B. finegoldii* | BaFIN-F | CCGGATGGCATAGGATTGTC | - | | 40) |
| | BaFIN-R | CGTAGGAGTTTGGACCGTGT | - | | |
| *B. fragilis* | FW3 | AGGATTCCGGTAAAGGATGG | - | | original |
| | RV3 | GTTCAGGCTAGCGCCCATT | - | | |
| *B. intestinalis* | BaINT-F | AGCATGACCTAGCAATAGGTTG | - | | 40) |
| | BaINT-R | ACGCATCCCCATCGATTAT | - | | |
| *B. plebeius* | BaPLE-F | ATCATTAAAGATTTATCGGTGTACG | - | | 40) |
| | BaPLE-R | ACTTTCACAGCTGACTTAACGAC | - | | |
| *B. stercoris* | B.ste-TaqMan-F | GCTTGCTTTGATGGATGGC | B.ste-TaqMan-Prb | CCAACCTGCCGACAACACTGGGATA | 39) |
| | B.ste-TaqMan-R | CATGCGGGAAAACTATGCC | - | | |
| *B. thetaiotaomicron* | B.the-TaqMan-F | GCAAACTGGAGATGGCGA | B.the-TaqMan-Pro | TCGATGGGGATGCGTTCCATTAGG | 39) |
| | B.the-TaqMan-R | AAGGTTTGGTGAGCCGTTA | - | | |
| *B. uniformis* | B.uni-TaqMan-F | TCTTCCGCATGGTAGAACTATTA | B.uni-TaqMan-Prb | CGTTCCATTAGGTTGTTGGCGGGG | 39) |
| | B.uni-TaqMan-R | ACCGTGTCTCAGTTCCAATGTG | - | | |
| *B. vulgatus* | B.vul-TaqMan-F | CGGGCTTAAATTGCAGATGA | B.vul-TaqMan-Prb | TGAAAGCCGTAAGCCGCAAGG | 39) |
| | B.vul-TaqMan-R | CATGCAGCACCTTCACAGAT | - | | |

The TaqMan probes were labeled with the fluorescent dyes 6-carboxyfluorescein (FAM) at the 5' end and 6-carboxytetramethylrhodamine (TAMRA) at the 3' end.

Supplementary Table 9. Composition of test foods (daily dose) using in human study.

| | Test food FL（mg） | Test food αCD（mg） | Placebo（mg） |
|---|---|---|---|
| Flaxseed lignan | 200.1 | 0.0 | 0.0 |
| Alpha-cyclodextrin | 0.0 | 200.1 | 0.0 |
| Maltitol | 487.1 | 487.1 | 687.2 |
| Tricalcium phosphate | 19.4 | 19.4 | 19.4 |
| Hydroxypropyl Cellulose | 9.0 | 9.0 | 9.0 |
| Calcium stearate | 30.0 | 30.0 | 30.0 |
| Starch | 3.0 | 3.0 | 3.0 |
| Silicon dioxide | 1.5 | 1.5 | 1.5 |
